# Discovery and characterization of a Gram-positive Pel polysaccharide biosynthetic gene cluster

**DOI:** 10.1101/768473

**Authors:** Gregory B. Whitfield, Lindsey S. Marmont, Cedoljub Bundalovic-Torma, Erum Razvi, Elyse J. Roach, Cezar M. Khursigara, John Parkinson, P. Lynne Howell

## Abstract

Our understanding of the biofilm matrix components utilized by Gram-positive bacteria, and the signalling pathways that regulate their production are largely unknown. In a companion study, we developed a computational pipeline for the unbiased identification of homologous bacterial operons and applied this algorithm to the analysis of synthase-dependent exopolysaccharide biosynthetic systems (https://doi.org/10.1101/769745). Here, we explore the finding that many species of Gram-positive bacteria have operons with similarity to the *Pseudomonas aeruginosa pel* locus. Our characterization of the *pelDEA_DA_FG* operon from *Bacillus cereus* ATCC 10987, presented herein, demonstrates that this locus is required for biofilm formation and produces a polysaccharide structurally similar to Pel. We show that the degenerate GGDEF domain of the *B. cereus* PelD ortholog binds cyclic-3’,5’-dimeric guanosine monophosphate (c-di-GMP), and that this binding is required for biofilm formation. Finally, we identify a diguanylate cyclase, CdgF, and a c-di-GMP phosphodiesterase, CdgE, that reciprocally regulate the production of Pel. The discovery of this novel c-di-GMP regulatory circuit significantly contributes to our limited understanding of c-di-GMP signalling in Gram-positive organisms. Furthermore, conservation of the core *pelDEA_DA_FG* locus amongst many species of Bacilli, Clostridia, Streptococci, and Actinobacteria suggests that Pel may be a common biofilm matrix component in many Gram-positive bacteria.

**Author summary:** The Pel polysaccharide is required for biofilm formation in *P. aeruginosa* and we have previously found that the genes necessary for biosynthesis of this polymer are broadly distributed across Gram-negative bacteria. Herein, we show that many species of Gram-positive bacteria also possess Pel biosynthetic genes and demonstrate that these genes are used *Bacillus cereus* for biofilm formation. We show that Pel production in *B. cereus* is regulated by c-di-GMP and have identified two enzymes, a diguanylate cyclase, CdgF, and a phosphodiesterase, CdgE, that control the levels of this bacterial signalling molecule. While Pel production in *B. cereus* also requires the binding of c-di-GMP to the receptor PelD, the divergence of this protein in Streptococci suggests a c-di-GMP independent mechanism of regulation is used in this species. The discovery of a Pel biosynthetic gene cluster in Gram-positive bacteria and our characterization of the *pel* operon in *B. cereus* suggests that Pel is a widespread biofilm component across all bacteria.

## Introduction

Bacteria are routinely challenged by a variety of environmental conditions that test their ability to survive. One mechanism employed by a wide range of microorganisms to withstand these challenges is to form a structured multicellular community, or biofilm. Bacteria growing as a biofilm are more tolerant of the presence of toxic compounds [1, 2] and predation by protists [3, 4], and exhibit increased persistence during infection due to antimicrobial tolerance [5, 6] and the ability to evade the host immune response [7, 8]. As a result, biofilm formation is linked not only to chronic bacterial infection in humans [9] [10], and plants and animals of economic importance [11–13], but also to the contamination of industrial facilities involved in processing food [14, 15], and pulp and paper [16].

Biofilm formation requires the production of extracellular material, or matrix, that allows bacteria to adhere to each other and/or to a surface [17]. While this matrix can be composed of protein adhesins, functional amyloid, extracellular DNA, and polysaccharides, the relative importance and abundance of each of these components varies greatly between bacterial species [17]. The complexity of the biofilm matrix is further amplified by the unique structural and functional characteristics of the matrix components produced by different bacteria, such as the cepacian polysaccharide produced by *Burkholderia* species [18], the CdrA adhesin of Pseudomonads [19], or TasA fibres from Bacilli [20]. Indeed, there is even variation in the utilization of matrix components for biofilm formation between different strains of the same species [21]. As a result of this complexity, identification of the biofilm matrix composition of many clinically and industrially relevant biofilm-forming organisms remains unresolved [22].

Despite the complexity of biofilm matrix structure, there are components that are utilized by a wide range of organisms. In particular, cellulose [23] and poly-β-(1,6)-*N*-acetyl-D-glucosamine (PNAG; [24]) have been identified as biofilm components across many bacterial genera. In a companion paper [25], we developed a computational pipeline that enables the unbiased identification of functionally related gene clusters from genome sequence data. Applying this pipeline to all bacterial phyla, we performed a systematic search for synthase-dependent polysaccharide biosynthetic operons involved in cellulose, PNAG, alginate and Pel polysaccharide production. From this search we identified *pel* gene clusters not only in Gram-negative species where it was originally identified but, unexpectantly, a range of Gram-positive species. Herein, we show that one of these *pel* gene clusters, *pelDEA_DA_FG,* from *Bacillus cereus* ATCC 10987, is involved in the biosynthesis of a Pel-like polysaccharide which is essential for biofilm formation by this organism. We identified a diguanylate cyclase, CdgF, and a cyclic-3’5’-dimeric guanosine monophosphate (c-di-GMP) phosphodiesterase, CdgE, that regulate biofilm formation and demonstrate that production of the polysaccharide is regulated post-translationally through binding of c-di-GMP to the inhibitory site of a degenerate GGDEF domain-containing receptor similar to PelD from *Pseudomonas aeruginosa* [26]. Our data not only expand the range of organisms in which Pel loci are found, but also identify novel, functionally distinct relatives of the Pel locus in Gram-positive bacteria that were missed in prior analyses [27]. This study establishes Pel as contributing to the arsenal of biofilm formation mechanisms acquired by Gram positives, as well as providing a rare example of post-translational c-di-GMP-mediated regulation of biofilm formation in this group of organisms.

## Results

### Identification of Pel biosynthetic loci in Gram-positive bacteria

The Pel polysaccharide was first identified in the Gram-negative opportunistic pathogen *P. aeruginosa*, where its biosynthesis has been linked to the *pelABCDEFG* operon [28]. Bioinformatic and biochemical analysis of the protein products of the *pel* operon has revealed a complex membrane-spanning molecular machine (Fig. S1). Previous work from our lab identified *pel* operons in bacterial genomes through bioinformatic searches using the outer membrane lipoprotein PelC as a query sequence. We used this small outer membrane lipoprotein as it is unique to Pel biosynthesis [27]. While this work expanded our knowledge of bacteria with *pel* operons considerably, the use of PelC as a search sequence limits this analysis to Gram-negative bacteria. To overcome this, we developed a computational pipeline that allows for the unbiased identification of homologous bacterial operons [25]. A search for *pel* loci using the protein coding sequences of the *P. aeruginosa* PAO1 *pel* operon as a reference set identified *pelF* and *pelG* loci in 43 Gram-positive phyla, including commonly studied species of Bacilli, Clostridia, and Streptococci [25]. Manual examination of the genomic context of the *pelFG* loci revealed the presence of genes whose protein products are consistent with the functions of PelA, PelD, and PelE when analyzed by BLAST [29] and Phyre^2^[30]. The complete *pel*-like *pelDEAFG* operon (Fig. 1) would provide the minimally required components for Pel polymerization and transport across the cytoplasmic membrane of Gram-positive bacteria. We used these genes as a basis to perform a second search of fully sequenced Gram-positive genomes that identified new loci that were missed during the initial screen, ultimately identifying 161 Gram-positive organisms harboring *pel-*like operons (Table S1; [25]).

**Figure 1:**
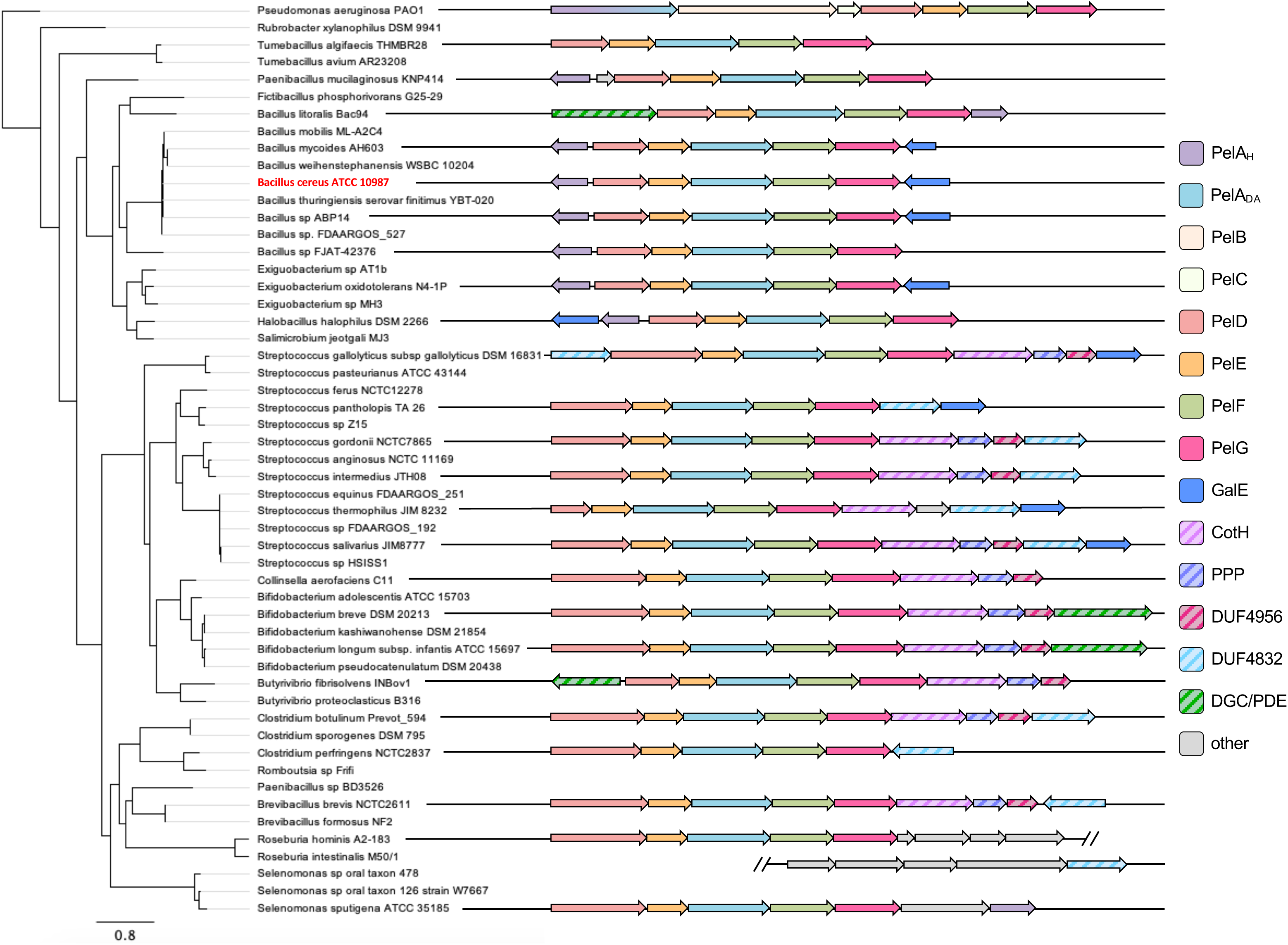
Representative Gram-positive *pel*-like operon architectures. (*left*) Phylogenetic tree generated from multiple sequence alignment of PelG protein sequences from select bacterial species identified through iterative hidden Markov model (HMM) searches. (*right*) Representative *pel*-like operon architectures from the corresponding organisms on the left. Open reading frames are represented as arrows, with the directionality of transcription indicated by the arrow direction. Open reading frames and overall operon architectures are drawn to scale. Arrow colours correspond to predicted protein functions, which are listed in the legend to the right. Open reading frames identified as ‘other’ correspond to genes that were not conserved amongst identified *pel*-like loci. *B. cereus* ATCC 10987 is highlighted in red. PelA_H_, PelA hydrolase-like domain; PelA_DA_, PelA deacetylase-like domain; PPP, poly-phosphate polymerase; DUF, domain of unknown function; DGC/PDE, diguanylate cyclase/phosphodiesterase-like domain.

While there existed remarkable similarity between the Gram-positive *pel*-like operon identified herein and the *pelABCDEFG* operon from *P. aeruginosa*, there were also several notable differences in protein domain architecture. First, Gram-positive orthologs of PelA have only a polysaccharide deacetylase domain (PelA_DA_), while *P. aeruginosa* PelA has both deacetylase and glycoside hydrolase domains (Fig. 1; [31]). Second, analysis of PelD-like proteins using Phyre^2^ revealed a number of significant genera-specific differences in their predicted domain architecture (Fig. S2). While the majority of Bacilli had PelD proteins with similar domain architectures and predicted functions to that of *P. aeruginosa* PelD [26], all the *Brevibacillus* strains identified had an additional predicted short-chain dehydrogenase/reductase (SDR) domain at their amino terminus (Fig. S2). Furthermore, analysis of potential co-factor binding and catalytic residues in this region suggests that this domain may be a functional epimerase, with strong similarity to UDP-glucose-4-epimerases. Strikingly, all other extended-length PelD proteins, including those identified in Streptococci, Clostridia, and Actinobacteria, also contained a predicted epimerase domain at the amino terminus (Fig. S2). However, in these bacteria the residues predicted to be required for co-factor binding and epimerase activity are significantly altered, suggesting that these domains are likely catalytically inactive and therefore performing some other function besides epimerization.

A further functional modification to PelD was identified specifically in Streptococci as these species lack the predicted degenerate GGDEF domain (Fig. S2). Streptococcal species are not known to use c-di-GMP as a signalling molecule and do not typically encode any DGC or PDE enzymes in their genomes [32, 33]. Therefore, it is perhaps not surprising that the domain of PelD necessary for c-di-GMP recognition has been lost in these species. In PelD from *S. thermophilus* JIM8232 there was a further modification not seen in any other PelD proteins, in which the putative amino-terminal epimerase domain was also absent, leaving just the transmembrane region and an adjoining GAF domain (Fig. S2). Analysis of the genomic region surrounding JIM8232 *pelD* revealed the presence of several transposase genes upstream of the *pelD* ORF, which may account for the loss of the amino-terminal epimerase region (Fig. S6).

With the exception of these differences in PelD domain architecture, the Gram-positive *pel*-like loci identified herein comprised a conserved set of genes whose sequence, predicted protein function, and synteny is exemplified by the *pelDEA_DA_FG* operon from *B. cereus* ATCC 10987 (Fig. 2A). Along with this core set of genes, we identified numerous accessory genes within close proximity to the *pelDEA_DA_FG* cluster that are irregularly distributed amongst the species identified (Fig. 1). Interestingly, the synteny of these accessory genes with respect to *pelDEA_DA_FG* cluster into two main groups. First, genes whose protein products exhibit similarity to the glycoside hydrolase domain of *P. aeruginosa* PelA [31], as well as the UDP-glucose-4-epimerase GalE [34] were commonly identified as divergently transcribed open reading frames (ORFs) flanking the *pelDEA_DA_FG* operon amongst Bacilli (Fig. S3). Orthologs of *galE* are also found at various positions within the *pel* locus of non-Bacilli (Fig. S3 and S4).

**Figure 2:**
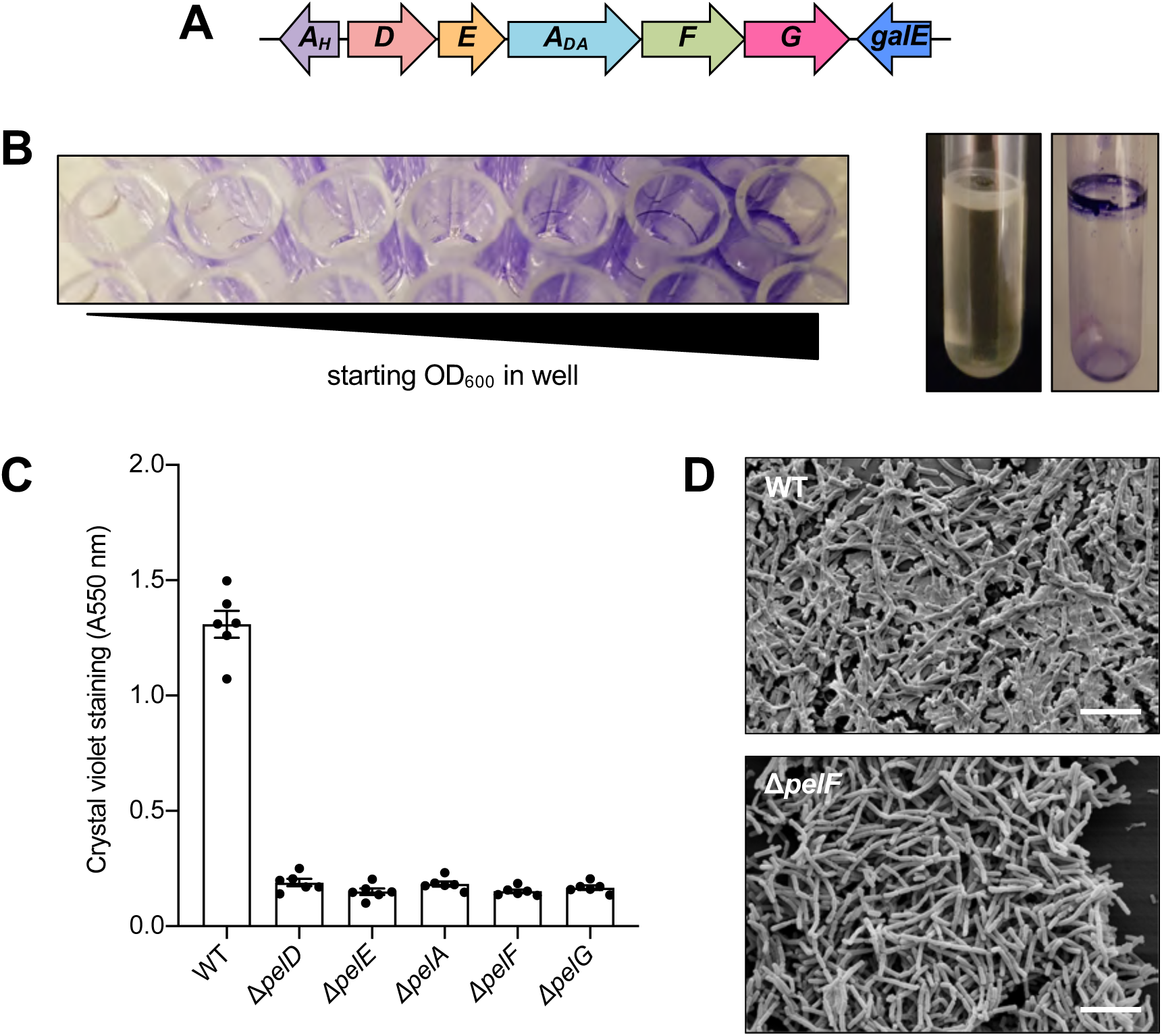
*B. cereus* ATCC 10987 forms biofilms that are dependent on the *pelDEA_DA_FG* operon. (A) *B. cereus* ATCC 10987 *pel* operon architecture. Open reading frames are represented as arrows, with the directionality of transcription indicated by the arrow direction. Open reading frames and the overall operon architecture is drawn to scale. Arrow colours correspond to predicted protein functions as defined in Fig. 1. (B) Air-liquid interface biofilm formation by ATCC 10987 in a 96-well microtitre plate (left) and a borosilicate glass tube (right). (C) Biofilm formation of the indicated strains of *B. cereus* ATCC 10987 assessed by the crystal violet assay. Error bars represent the standard error of the mean of six independent trials. (D) Scanning electron micrographs of the indicated strains of *B. cereus* ATCC 10987. Scale bars = 10 μm.

The second accessory gene group consisted of a contiguous set of genes whose protein products are similar to the kinase CotH [35], a poly-phosphate polymerase, and two conserved domains of unknown function, DUF4956 and DUF4832, that are contiguous with the *pelDEA_DA_FG* gene cluster in Streptococcal and Clostridial species (Fig. S4). A variant of this architecture is found in *Streptococcus gallolyticus* strains, where the ORF predicted to encode the DUF4832-containing protein is located upstream of, and contiguous with, *pelDEA_DA_FG* (Fig. S4). In Bifidobacterial species, the DUF4832 ORF is replaced by a gene predicted to encode a dual diguanylate cyclase/phosphodiesterase (Fig. S5).

Some species exhibit noteworthy variations in the predicted function and arrangement of the accessory genes (Fig. 1). In *Streptococcus thermophilus*, there are significant variations in operon architectures from one strain to the next, some of which are interrupted by transposons (Fig. S6). In *Roseburia* spp., the *pelDEA_DA_FG* locus is part of a larger putative gene cluster that contains genes whose functions are consistent with capsule or teichoic acid biosynthesis (Fig. S7). It remains unclear whether these operons are functional given these variations, or whether they are involved in the production of a polymer unrelated to Pel.

While the array of Gram-positive species containing the putative *pelDEA_DA_FG* locus spanned a broad phylogenetic distribution (Fig. 1), several genera are noticeably absent, including Listeria and Staphylococci. However, 53 Staphylococcal genomes, largely comprising strains of the human pathogen *Staphylococcus aureus*, were found from our accompanying bioinformatics analysis to possess operons involved in the production of the extracellular polysaccharide PNAG [25], which likely reflects its key role in virulence [36]. Collectively, the data presented here suggests that many Gram-positive bacteria have a previously unrecognized genetic locus that, at its core, is remarkably similar to the *pelABCDEFG* operon from *P. aeruginosa*.

### *B. cereus* ATCC 10987 utilizes the *pel*-like operon for biofilm formation

Next, we wanted to determine whether these operons were used for biofilm formation. We reasoned that the relatively simple architecture of the *pel*-like locus in Bacilli may form the minimal genetic unit required for Pel biosynthesis in Gram-positive bacteria, and as ATCC 10987 has previously been reported to produce a significant amount of biofilm biomass [37–39] we selected this strain as a representative model to determine whether the *pel-*like locus was functional (Fig. 2A). We were able to replicate the pellicle phenotype associated with Pel production in *P. aeruginosa* using routine biofilm assays (Fig. 2B; [28]). To determine whether the *pel*-like locus (*BCE_5583* – *BCE_5587*) contributes to biofilm formation in ATCC 10987 and which genes are essential for this process, we generated markerless deletions of each gene. We then compared the ability of each deletion mutant to form biofilm using the crystal violet microtitre plate assay (Fig. 2C). We found that when any of *pelDEA_DA_FG* were deleted, adhesion to the wells of the plastic microtitre dish was significantly reduced (Fig. 2C).

To assess whether these results may be due to polar effects resulting from the engineered chromosomal gene deletions, each mutant was complemented with a copy of the deleted gene on a xylose-inducible replicating plasmid, pAD123-P_xyl_, or the equivalent empty vector as a vehicle control. The complemented deletion mutants, but not those harbouring the empty vector control, restored biofilm formation at the air-liquid interface and adherence to plastic (Fig. S8), suggesting that the *pelDEA_DA_FG* operon is utilized for adherence of ATCC 10987 to chemically distinct surfaces and that loss of this phenotype is not due to polar effects from the allelic replacement process.

We hypothesized that the *pelDEA_DA_FG* operon from ATCC 10987 may be involved in the production of an extracellular matrix material, similar to the function of the *pelABCDEFG* operon from *P. aeruginosa* [28]. To examine matrix formation at a microscopic level, we grew wild-type ATCC 10987 or the isogenic Δ*pelF* mutant on plastic coverslips and analyzed the adherent bacteria by scanning electron microscopy (SEM; Fig. 2D). These images showed wild-type ATCC 10987 surrounded by, and embedded within, copious amounts of extracellular material (Fig. 2D, top). In contrast, the Δ*pelF* mutant produced little-to-no visible extracellular material (Fig. 2D, bottom). Combined, these results suggest that each of *pelD*, *pelE*, *pelA_DA_*, *pelF*, and *pelG* are necessary for biofilm formation (Fig. 2C), and that the polymer produced is the dominant matrix component produced by this strain of *B. cereus* under the conditions tested.

### The *pelDEA_DA_FG* operon is involved in the production of a Pel-like polysaccharide

To determine whether the extracellular material produced by the *pelDEA_DA_FG* operon was compositionally similar to Pel produced by *P. aeruginosa*, we utilized the *Wisteria floribunda* lectin (WFL), which recognizes terminal *N*-acetyl-galactosamine (GalNAc) moieties [40] and has previously been shown to specifically bind the *P. aeruginosa* Pel polysaccharide [41]. We analyzed the ability of WFL to bind to the biomass produced by wild-type ATCC 10987 or the isogenic Δ*pelF* mutant by fluorescence microscopy (Fig. 3A). As a control, a *P. aeruginosa* strain engineered to overexpress the Pel polysaccharide, PAO1 Δ*wspF* Δ*psl* P_BAD_*pel* [31], and its isogenic Δ*pelF* mutant, were also analyzed (Fig. 3A). Under the conditions of this experiment, both wild-type *B. cereus* and *P. aeruginosa* strains were able to form a confluent layer of cells covering a significant portion of the field of view, while the Δ*pelF* mutants were observed as smaller aggregates or individual cells (Fig. 3A, brightfield panels), consistent with the ability of these strains to form biofilms that are dependent on *pelF* (Fig. 2C; [28]). To detect biofilm biomass, 4’,6-diamidino-2-phenylindole (DAPI) was used as a marker for DNA, which would be present within intact cells as well as in the biofilm matrix as extracellular DNA (eDNA). This showed that DAPI-stained biomass co-localized with cells observed in brightfield microscopy (Fig. 3A, DAPI panels, compare to brightfield panels). Finally, analysis of WFL-Texas Red fluorescence revealed co-localization of WFL with biomass detected by DAPI staining in the *B. cereus* and *P. aeruginosa* wild-type strains, while no co-localization was observed in the *pelF* mutants (Fig. 3A).

**Figure 3:**
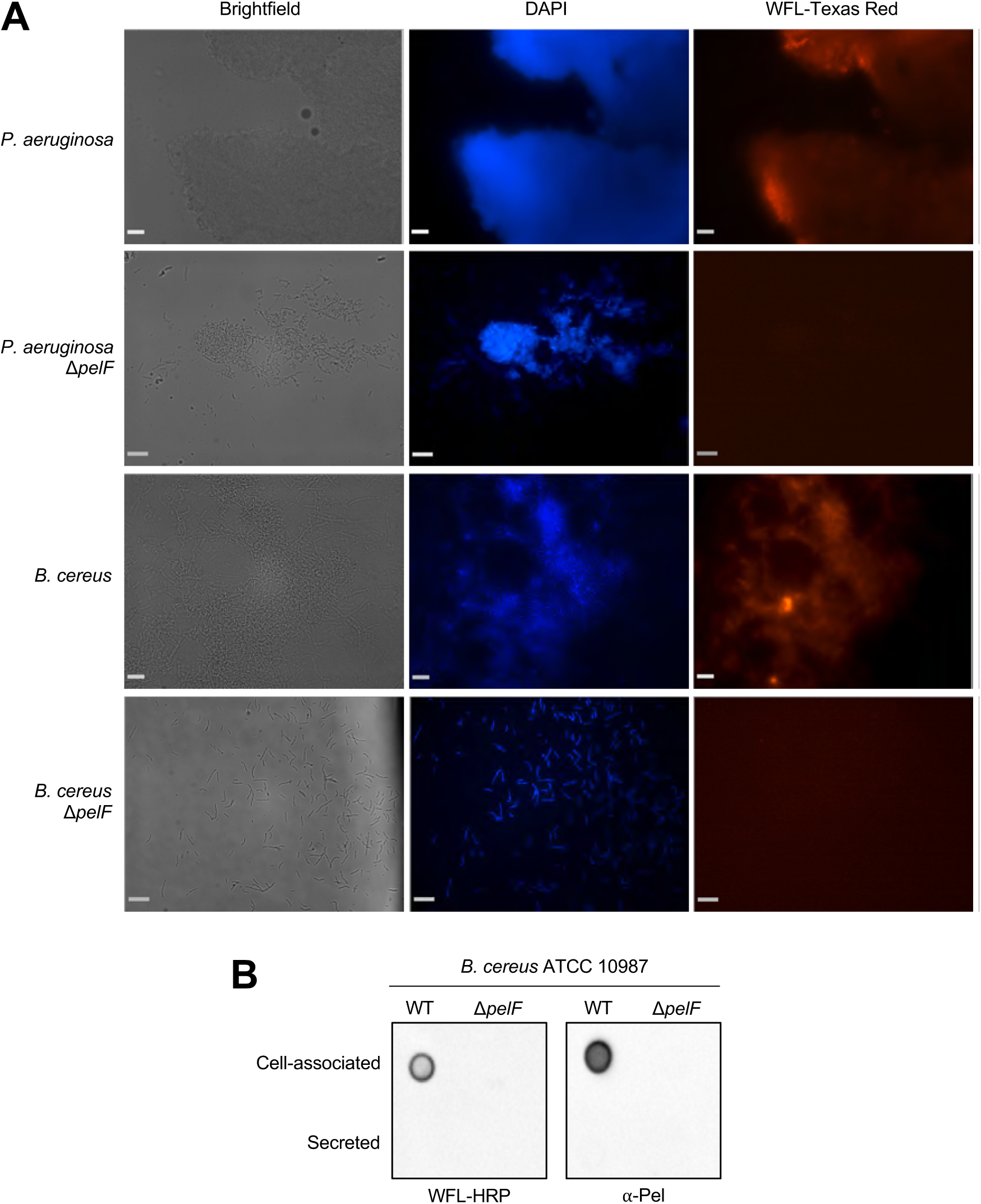
The *pel* operon of *B. cereus* ATCC 10987 is involved in the generation of a Pel-like extracellular material. (A) Brightfield and epifluorescence microscopy of *P. aeruginosa* (PAO1 Δ*wspF* Δ*psl* P_BAD_*pel*) wild-type and Δ*pelF* mutant, and *B. cereus* ATCC 10987 wild-type and Δ*pelF* mutant biofilms grown for 24 h on poly-L-lysine coated glass coverslips and stained with DAPI, to detect DNA, and Wisteria floribunda lectin (WFL) conjugated to Texas red to detect Pel-like extracellular material. Scale bar = 250 μm. (B) Dot blot of extracellular material from *B. cereus* ATCC 10987 or the Δ*pelF* mutant grown for 24 h at 30 °C with shaking, separated into crude cell-associated and secreted fractions. Extracellular material was detected using WFL conjugated to horseradish peroxidase (HRP; *left*) or 12-Pel primary antibody with HRP-conjugated secondary antibody (*right*).

To further analyze the extracellular material produced by *B. cereus*, cell-associated and secreted fractions from liquid cultures of ATCC 10987 or the Δ*pelF* mutant were analyzed by dot blot using either WFL or a Pel-specific antibody (Fig. 3B; [31]). Both WFL and the α-Pel antibody recognized cell-associated material from wild-type ATCC 10987, but not the Δ*pelF* mutant (Fig. 3B). Collectively, these results suggest that the extracellular material produced by the *pelDEA_DA_FG* operon of ATCC 10987 is predominantly carbohydrate in nature, contains terminal GalNAc moieties, and has a chemical composition similar to that of the Pel polysaccharide from *P. aeruginosa* [41].

### *BCE_5582* is a glycoside hydrolase that influences ATCC 10987 biofilm formation

In addition to the core *pelDEA_DA_FG* operon identified in ATCC 10987, there were also two divergently transcribed accessory genes, *BCE_5582* and *BCE_5588*, upstream and downstream of the *pel* operon, respectively (Fig. S3). BCE_5588 is homologous to the UDP-glucose-4-epimerase, GalE, and a transposon screen published during the course of this study has already implicated this gene in biofilm formation [42]. As BCE_5582 was predicted by the Phyre^2^ algorithm [30] to adopt a structure similar to the endo-α-1,4-*N*-acetylgalactosaminidase domain of *P. aeruginosa* PelA (PelA_H_*^Pa^*; [43]), we refer to BCE_5582 as PelA_H_*^Bc^* herein. Since PelA encoded in the *pelDEA_DA_FG* operon is predicted to contain only a polysaccharide deacetylase domain (PelA_DA_), and we found orthologs of *pelA_H_*-like genes within or adjacent to many of the *pel* operons (Fig. 1, S3, S4), its function may be relevant to the biofilm formation phenotypes observed in ATCC 10987. To assess this hypothesis, a deletion mutant of *pelA_H_^Bc^* was generated and evaluated for biofilm formation using the crystal violet assay (Fig. 4A). Deletion of *pelA_H_^Bc^* led to a significant increase in adherent biomass versus wild-type ATCC 10987. To confirm that this phenotype was attributable to *pelA_H_^Bc^,* rather than the result of a lost negative regulatory element upstream of the *pelDEA_DA_FG* operon, this mutant was complemented with *pelA_H_^Bc^*on pAD123-P_xyl_. Complementation of the *pelA_H_^Bc^*mutant revealed that the hyper-biofilm phenotype of the mutant was due specifically to *pelA_H_^Bc^* (Fig. 4A).

**Figure 4:**
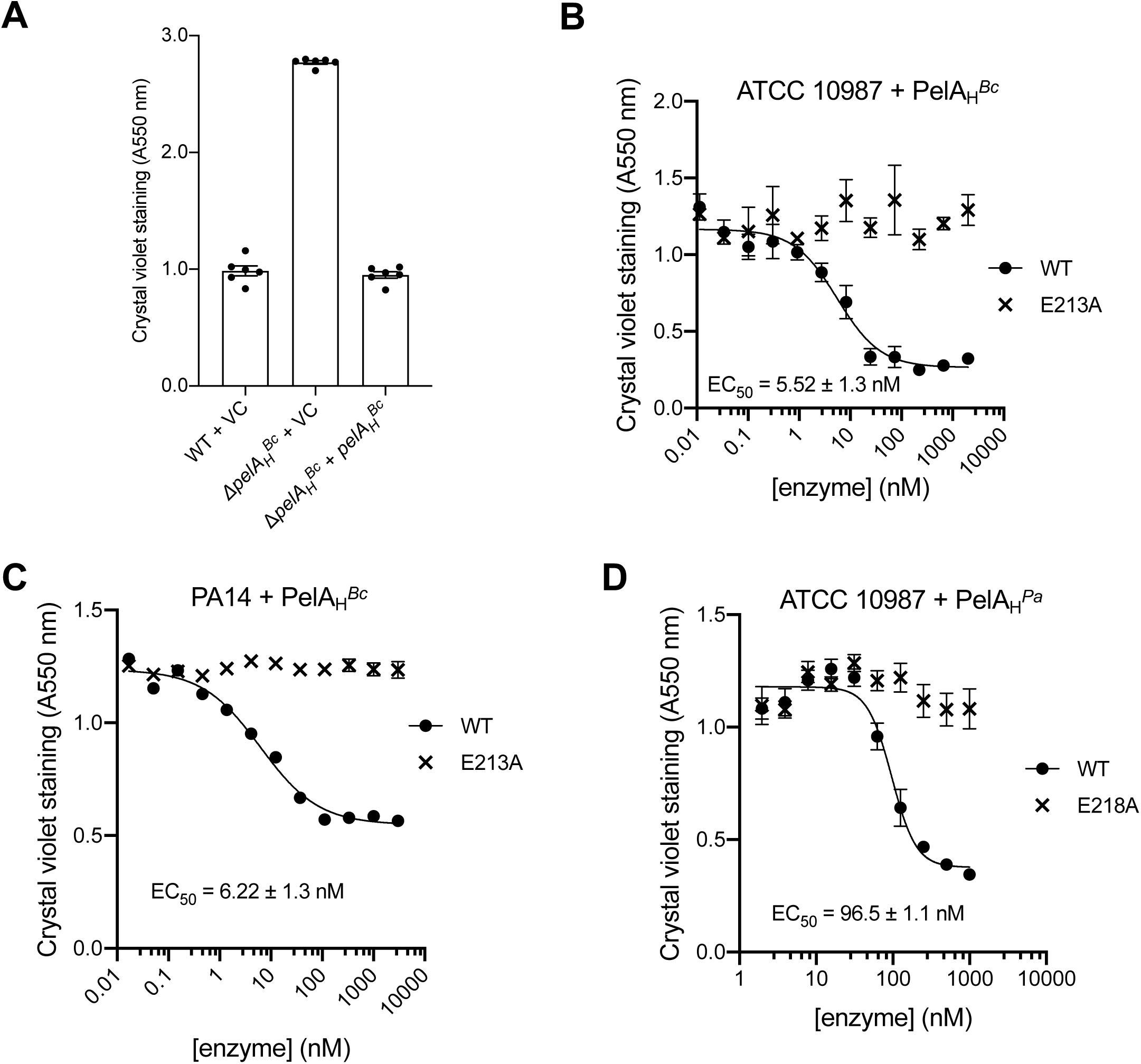
PelA_H_^*Bc*^ (BCE_5582) is a predicted glycoside hydrolase that is capable of disrupting pre-formed Pel-dependent biofilms. (A) Biofilm formation of the indicated strains of *B. cereus* ATCC 10987 assessed by the crystal violet assay. VC, empty vector control. (B-D) Dose-response curves generated by the exogenous application of the indicated glycoside hydrolase enzyme to pre-formed biofilms of the indicated strains. (B) Application of exogenous PelA_H_*^Bc^*, or the catalytic point variant E213A, to pre-formed ATCC 10987 biofilms. (C) Application of exogenous PelA_H_*^Bc^*, or the catalytic point variant E213A, to pre-formed *P. aeruginosa* PA14 biofilms. (D) Application of exogenous *P. aeruginosa* hydrolase PelA (PelA_H_*^Pa^*), or the catalytic point variant E218A, to pre-formed ATCC 10987 biofilms. Error bars represent the standard error of the mean of six independent trials. EC_50_ values were calculated using nonlinear least-squares fitting to a dose-response model.

As our data suggested that *pelA_H_^Bc^* may be playing a regulatory role in ATCC 10987 biofilm formation, and PelA_H_*^Bc^*is predicted to be a glycoside hydrolase, we sought to validate the function of PelA_H_*^Bc^*. Full-length PelA_H_*^Bc^*minus its signal sequence, residues 26-276, was therefore heterologously expressed and purified. Using an indirect *in vitro* assay for glycoside hydrolase activity [44, 45] we found that PelA_H_*^Bc^* was able to disrupt pre-formed ATCC 10987 biofilms in a concentration-dependent manner (EC_50_ = 5.52 ± 1.3 nM; Fig. 4B). To verify that this result was due to the activity of PelA_H_*^Bc^*, a residue predicted to be important for glycoside hydrolase catalytic activity, Glu 213, was mutated to alanine. When pre-formed biofilms were challenged with the E213A mutant, no disruption was observed (Fig. 4B). These results suggest that PelA_H_*^Bc^* is a glycoside hydrolase that is active against an extracellular polysaccharide produced by ATCC 10987, likely the product of the *pelDEA_DA_FG* locus.

Since glycoside hydrolases are specific for particular polysaccharide structures [46, 47], we reasoned that we could utilize PelA_H_*^Bc^*as a tool to further validate our hypothesis that the polysaccharide produced by ATCC 10987 is structurally similar to *P. aeruginosa* Pel [41]. We found that PelA_H_*^Bc^* disrupted pre-formed biofilms produced by the Pel-dependent *P. aeruginosa* strain PA14 [21] in a concentration-dependent manner (EC_50_ = 6.22 ± 1.3 nM; Fig. 4C). As with ATCC 10987, the PelA_H_*^Bc^*E213A mutant had no effect on PA14 biofilms (Fig. 4C). Since the WFL binding assay demonstrated that the polymers produced by *B. cereus* and *P. aeruginosa* are comparable (Fig. 3), we used the endo-α-1,4-*N*-acetylgalactosaminidase PelA_H_*^Pa^* [43] in a reciprocal experiment to challenge pre-formed ATCC 10987 biofilms. This led to a concentration-dependent decrease in biofilm formation (EC_50_ = 96.5 ± 1.1 nM; Fig. 4D), while a catalytically inactive mutant of PelA_H_*^Pa^*had no effect on pre-formed ATCC 10987 biofilms (Fig. 4D). This data, along with binding of the GalNAc-specific WFL lectin [40] and the Pel-specific antibody (Fig. 3; [31]), supports the hypothesis that the biosynthetic product of the ATCC 10987 *pel* locus is structurally similar to *P. aeruginosa* Pel, including the presence of α-1,4-linked GalNAc moieties.

### Binding of c-di-GMP to ATCC 10987 PelD is required for biofilm formation

In *P. aeruginosa*, Pel biosynthesis is regulated post-translationally through binding of c-di-GMP to the inhibitory site of the degenerate GGDEF domain of PelD (PelD*^Pa^*; [26, 48]). As the *B. cereus* ATCC 10987 *pel* operon contains a gene with predicted similarity to PelD*^Pa^* (Fig. 2A), we next investigated whether biofilm formation in *B. cereus* is post-translationally regulated using a similar mechanism. To better assess the function of *B. cereus* PelD (PelD*^Bc^*), a structural model of its GAF and GGDEF domains was generated and the residues from PelD*^Pa^*known to be crucial for c-di-GMP binding were mapped onto the PelD*^Bc^* structure (Fig. 5A). As the presence of an intact RxxD motif in PelD*^Bc^* suggested that it may be capable of binding c-di-GMP in a similar manner to that of PelD*^Pa^*, we expressed and purified the cytosolic domain of PelD*^Bc^* (PelD*^Bc^*_170-403_) and analyzed the ability of this fragment to bind c-di-GMP using isothermal titration calorimetry (ITC; Fig. 5B, left). This revealed that PelD*^Bc^*_170-403_ was capable of binding c-di-GMP with a K_D_ of 103 ± 14 nM under the conditions tested. Residues predicted to be necessary for the binding of c-di-GMP, including R363 and D366 from the RxxD motif and R395 (Fig. 5A), were mutated to alanine in the PelD*^Bc^*_170-403_ construct and c-di-GMP binding was again examined by ITC (Fig. 5B). We could not detect any heats of emission or absorption for the alanine point variants when c-di-GMP was titrated into the protein solution, suggesting that these residues are important for c-di-GMP binding to PelD*^Bc^*. To test whether binding of c-di-GMP to PelD*^Bc^* is necessary for the biosynthesis of Pel as previously demonstrated for PelD*^Pa^*[26], the R363A, D366A, and R395A point mutations were introduced onto the chromosome of ATCC 10987 and biofilm formation was assessed by the crystal violet assay (Fig. 5C). Strains of ATCC 10987 harbouring R363A, D366A, or R395A mutations in PelD were unable to adhere to the plastic walls of a 96-well microtitre plate, similar to the results obtained for the Δ*pelF* mutant (Fig. 5C). Taken together our *in vivo* and *in vitro* analysis suggests that binding of c-di-GMP to PelD is necessary for biofilm formation in ATCC 10987 and that, like PelD*^Pa^*, PelD*^Bc^* likely binds c-di-GMP at its I-site using the RxxD motif [26].

**Figure 5:**
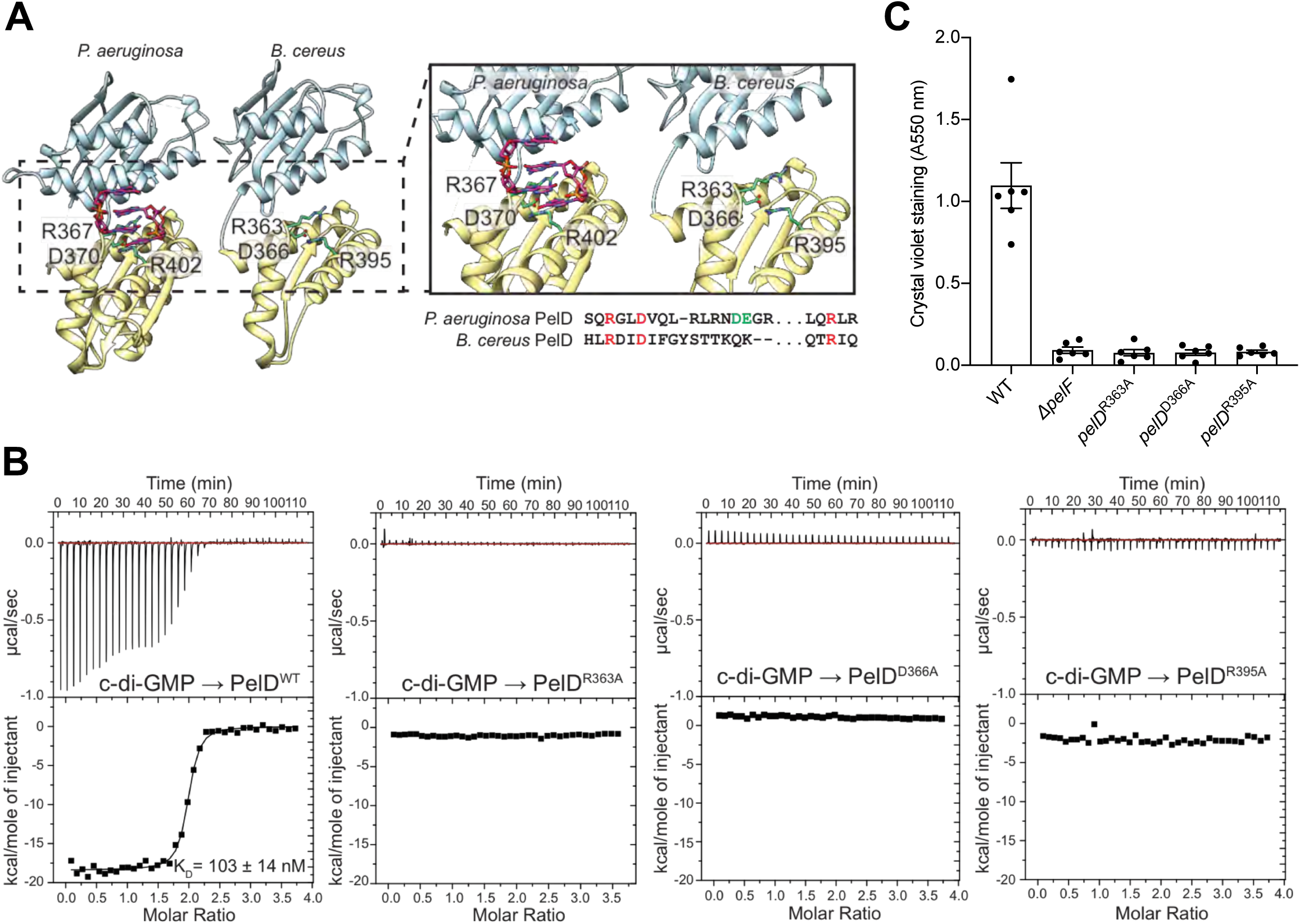
Binding of c-di-GMP to PelD is required for *B. cereus* ATCC 10987 biofilm formation. (A) Structure of *P. aeruginosa* PelD_156-455_ (PDB code: 4DN0), and model of *B. cereus* PelD_170-403_. Inset: close up of the I-site and residues involved in the binding of c-di-GMP. The equivalent *B. cereus* residues to Arg367, Asp370, and Arg402 of *P. aeruginosa* were identified as Arg363, Asp366, and Arg395, and are represented as green sticks for clarity. The sequence containing these residues is indicated below the inset. (B) Isothermal titration calorimetry experiments for the wild-type PelD*^Bc^*_150-407_ and the indicated mutants. Each experiment displays the heats of injection (top panel) and the normalized integration data as a function of the molar syringe and cell concentrations (bottom panel). The calculated dissociation constant (K_D_) for the wild-type protein is indicated on the figure. (C) Biofilm formation of the indicated strains of *B. cereus* ATCC 10987 assessed by the crystal violet assay. Error bars represent the standard error of the mean of six independent trials.

### Biofilm formation by ATCC 10987 is regulated by the diguanylate cyclase CdgF and the phosphodiesterase CdgE

All bacterial species studied to date, including *P. aeruginosa*, regulate intracellular levels of c-di-GMP through the reciprocal activities of c-di-GMP synthesizing diguanylate cyclases (DGCs) and c-di-GMP degrading phosphodiesterases (PDEs; [49]). DGCs and PDEs contain GGDEF, and EAL or HD-GYP domains, respectively. In *P. aeruginosa*, there are over 40 DGCs and PDEs with overlapping regulatory control over various c-di-GMP dependent phenotypes [50], making it difficult to disentangle the contribution of specific c-di-GMP metabolic enzymes to measurable phenotypes. In a recent computational survey of 90 sequenced *B. cereus* group bacterial genomes, only 13 genes, termed *cdgA-M*, whose protein products were predicted to contain GGDEF, EAL, or HD-GYP domains, were identified [51]. Of these 13 potential c-di-GMP metabolic enzymes, CdgC, CdgJ, and CdgL contain degenerate sequence motifs and are thus unlikely to be enzymatically active, while CdgG and CdgM are narrowly distributed amongst the genomes analyzed and absent from ATCC 10987 [51]. Based on these results, we hypothesized that the c-di-GMP regulatory network in ATCC 10987 may only encompass eight c-di-GMP metabolic enzymes, CdgA, CdgB, CdgD, CdgE, CdgF, CdgH, CdgI, and CdgK. Analyzing the results of the deletion and overexpression of each c-di-GMP metabolic gene in *Bacillus thuringiensis* 407, a species closely related to *B. cereus*, revealed that two genes, *cdgE* and *cdgF*, had dominant yet opposing effects on intracellular c-di-GMP concentration and biofilm formation levels. While both CdgE and CdgF are predicted to have functional GGDEF and EAL domains, DGC activity was clearly associated with CdgF and PDE activity with CdgE [51].

To assess whether *cdgF* and *cdgE* play a role in biofilm formation by ATCC 10987, we generated *cdgF* and *cdgE* deletion mutant strains and analyzed biofilm formation using the crystal violet assay (Fig. 6). Deletion of *cdgF* in ATCC 10987 reduced adherence levels to that of a Δ*pelF* mutant (Fig. 6A), while complementation of this mutant with *cdgF* in pAD123-P_xyl_ led to restoration of adherence to wild-type levels at lower induction levels, and hyper-adherence at higher induction levels (Fig. 6A). Deletion of *cdgE* led to a hyper-adherence phenotype relative to the wild-type strain in the crystal violet assay, while overexpression of *cdgE* from pAD123-P_xyl_ reduced adherence in an inducer concentration-dependent manner, although adherence levels could not be restored to those observed for the wild-type strain (Fig. 6B). We hypothesize that this is due to difficulties overexpressing CdgE, which is predicted to contain ten transmembrane helices and would thus likely localize to the cytoplasmic membrane of ATCC 10987, versus the predicted cytosolic localization of CdgF [51]. To assess whether the increase and decrease in biofilm formation of the Δ*cdgE* and Δ*cdgF* mutants, respectively, were due specifically to production of Pel, dot blots of cell-associated and secreted extracellular material from liquid cultures of ATCC 10987 or the isogenic Δ*pelF*, Δ*cdgE*, or Δ*cdgF* mutants were analyzed by WFL and the Pel-specific antibody (Fig. 6C). This revealed that deletion of *cdgE* led to increased binding of both WFL and the α-Pel antibody versus the wild-type strain, with material appearing in the secreted fraction of the Δ*cdgE* mutant that was not present in the wild-type (Fig. 6C). In contrast, no signal was detected from the Δ*cdgF* mutant using WFL or the α-Pel antibody, similar to the Δ*pelF* mutant (Fig. 6C). This suggests that CgdE and CdgF play a significant role in the reciprocal regulation of Pel polysaccharide production from *B. cereus*. Thus, this marks the identification of a minimal c-di-GMP signalling network in *B. cereus* ATCC 10987 composed of the DGC CdgF, the PDE CdgE, and the receptor PelD, which together mediate biofilm formation through regulation of the production of a polysaccharide with significant similarity to Pel from *P. aeruginosa*.

**Figure 6:**
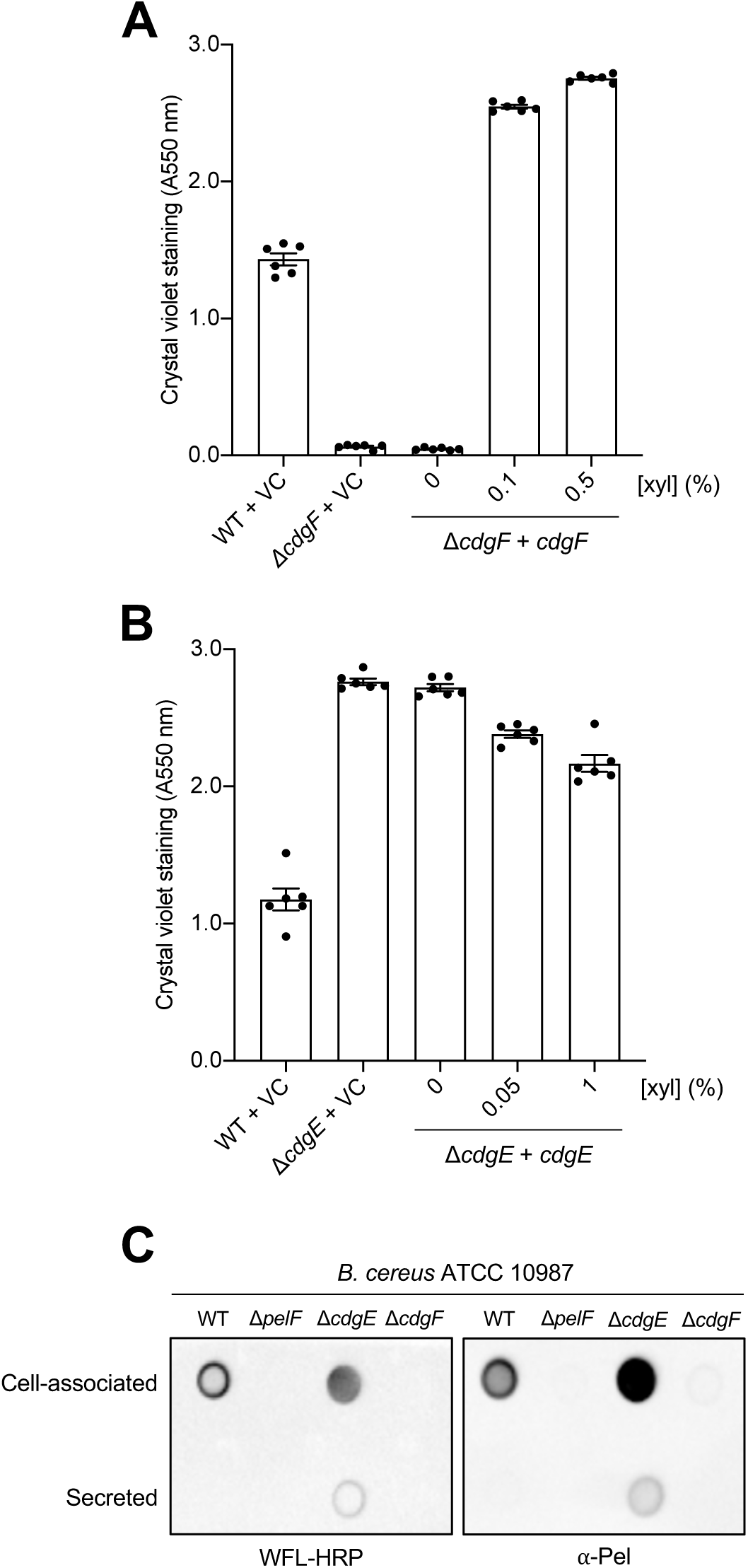
Biofilm formation by *B. cereus* ATCC 10987 is regulated by the diguanylate cyclase CdgF and the phosphodiesterase CdgE. (A) Biofilm formation by *cdgF* deletion and complementation strains assessed by the crystal violet assay. (B) Biofilm formation by *cdgE* deletion and complementation strains assessed by the crystal violet assay. VC, empty vector control; xyl, xylose. Error bars represent the standard error of the mean of six independent trials. (C) Dot blot of extracellular material from *B. cereus* ATCC 10987 or the isogenic Δ*pelF*, Δ*cdgE*, or Δ*cdgF* mutants grown for 24 h at 30 °C with shaking, separated into crude cell-associated and secreted fractions. Extracellular material was detected using WFL conjugated to horseradish peroxidase (HRP; *left*) or 12-Pel primary antibody with HRP-conjugated secondary antibody (*right*).

## Discussion

Identification of *pel* operons and the production of the Pel polysaccharide have not been described previously for any Gram-positive organism. Our computational pipeline predicted *pel* loci in 161 Gram-positive species (Table S1; [25]). Characterization of the *pelDEA_DA_FG* operon from *B. cereus* ATCC 10987 has validated these findings and shown that this strain uses the Pel polysaccharide for biofilm formation and that polymer production is regulated post-translationally by the binding of c-di-GMP to PelD. Furthermore, we show that Pel biosynthesis is reciprocally regulated by the DGC CdgF, and the PDE CdgE.

During our characterization of the *pelDEA_DA_FG* operon from *B. cereus* ATCC 10987, Okshevsky *et al.* published the results of a transposon mutant screen that identified genes required for biofilm formation in ATCC 10987 [42]. Transposon insertions in *BCE_5585*, *BCE_5586*, and *BCE_5587,* which we have termed *pelA_DA_*, *pelF*, and *pelG* herein, led to an impairment in biofilm formation and based on these results, Okshevsky *et al.* hypothesized that the *BCE_5583* – *BCE_5587* operon (herein *pelDEA_DA_FG*) may be involved in the production of an extracellular polysaccharide [42]. Two distinct transposon insertions were also observed in *cdgF*, which was the only DGC identified in the screen, as well as an insertion in *BCE_5588* that completely abrogated biofilm formation in ATCC 10987 [42]. *BCE_5588,* downstream of the *pel* operon, is similar to the UDP-glucose-4-epimerase GalE [34] and we hypothesized that BCE_5588 may be involved in the generation of the precursor nucleotide-sugars required for polysaccharide biosynthesis. The transposon insertion in *BCE_5588* generated by Okshevsky *et al.* confirms that this gene is required for biofilm formation in ATCC 10987. As this screen focused on mutants with impaired biofilm formation, genes whose deletion leads to hyper-biofilm formation, such as *BCE_5582* (*pelA_H_*) and *cdgE*, were not identified.

While it is tempting to speculate that the *pelDEA_DA_FG* operon identified in ATCC 10987 is a crucial locus for biofilm formation in *B. cereus*, our analysis only identified *pelDEA_DA_FG* operons in ∼17% of sequenced *B. cereus* genomes. For example, the locus is not present in the type strain ATCC 14579. This suggests that there may be significant strain-to-strain variability in matrix component utilization among *B. cereus* isolates. Indeed, eDNA, exopolysaccharide, and protein components have all been described as matrix components in various *B. cereus* strains [52–54]. As is the case amongst a diverse array of biofilm forming bacteria, eDNA has been shown to be crucial for biofilm formation in ATCC 14579 [52], the environmental *B. cereus* isolate AR156 [55], and ATCC 10987 [42]. First characterized in *Bacillus subtilis* [56], the exopolysaccharide producing *epsA-O* operon is also present in *B. cereus*, although its contribution to biofilm formation appears to be strain dependent. In the plant associated *B. cereus* strain 905, deletion of the *eps* locus does not affect biofilm formation [57], while in ATCC 10987 *eps* inactivation completely abrogated biofilm formation [42]. Strain-dependent utilization of different polysaccharide biosynthetic loci for biofilm formation has been observed in *P. aeruginosa*, where Pel and/or the Psl polysaccharide are variably required [21]. Finally, *B. cereus* also has two orthologs of the extracellular fibril forming *B. subtilis* TasA protein [58], TasA and CalY, which are required for biofilm formation in ATCC 14579 [54] but are only required for biofilm formation in strain 905 under certain culture conditions [57]. Whether ATCC 10987 utilizes TasA and/or CalY for biofilm formation is currently unknown.

Our bioinformatics analysis and validation of Pel production in ATCC 10987 described herein suggests that the *pel* operon is present and potentially functional in a variety of Gram-positive organisms. This is further validated by the work of Couvigny *et al*, who identified the gene we have called *pelG* as a requirement for biofilm formation by *Streptococcus thermophilus* and *Streptococcus salivarius* through a transposon mutagenesis screen [59]. Interestingly, the authors noted that 75% of *S. thermophilus* genomes (9 of 12) available at the time of their analysis did not contain the locus which we have attributed to Pel production, and suggested that this operon may be a remnant of commensal life that has been lost due to the adaptation of *S. thermophilus* to domestication and its use as a fermentation starter in the dairy industry [59]. Similarly, we note that this locus is only present in ∼26% (9 of 34) fully-sequenced *S. thermophilus* genomes available as of this writing. While *S. salivarius* is a commensal bacterium associated with the human gut and thus would be thought to more uniformly contain the *pel* locus as per the hypothesis of Couvigny *et al.*, we found that only 50% (5 of 10) fully-sequenced *S. salivarius* genomes contained the *pel* operon. Partial distribution *of pel* operons was also observed amongst *B. cereus* strains, where only 17% (14 of 79) fully-sequenced genomes contained the *pel* locus. The variable distribution of *pel* loci amongst *B. cereus*, *S. thermophilus*, and *S. salivarius* strains may represent strain-to-strain variability in matrix utilization and biofilm formation, which is supported by the variable biofilm formation capabilities of *S. thermophilus* isolates [59]. In contrast to this, we found that 100% (40 of 40) *Bifidobacterium breve* strains with fully-sequenced genomes contained a *pel* operon. However, given the scarcity of experimental data regarding biofilm formation by *Bifidobacterium breve* generally, we are unable to speculate as to why the *pel* locus is so well conserved in this species.

The motile to sessile transition in bacteria is regulated largely by c-di-GMP. To date, the mechanisms of signal regulation, signal transduction, and the phenotypic consequences of c-di-GMP signalling have largely been explored in Gram-negative bacteria [49]. This may be due in part to the extensive repertoire of c-di-GMP metabolic proteins, c-di-GMP receptors, and response elements present in model Gram-negative organisms such as *P. aeruginosa* [50], versus the comparably limited set of such proteins in model Gram-positive species such as *B. subtilis* and *Streptomyces coelicolor*, or their complete absence in Staphylococcal and Streptococcal species [33]. C-di-GMP signalling in *B. cereus* has remained a mystery until recently, when Fagerlund *et al.* published the first account of c-di-GMP metabolic enzymes in the *B. cereus* group [51]. Herein, we have shown that the putative DGC CdgF and the PDE CdgE play significant roles in the regulation of biofilm formation by ATCC 10987, and that this regulation is achieved, at least in part, through binding to the c-di-GMP receptor PelD. Our results suggest that PelD*^Bc^* contains a degenerate GGDEF domain that binds c-di-GMP via its intact I-site RxxD motif, which is similar to the mechanism used by *P. aeruginosa* PelD [26].

It is interesting to note that the analysis of the *B. cereus* group genomes by Fagerlund *et al*. did not identify PelD as a candidate GGDEF domain containing protein [51]. Furthermore, although *P. aeruginosa* PelD and *B. cereus* PelD are 33% identical across their entire length, algorithms that assess protein function based on primary sequence such as BLAST [29] and HMMER [60] are unable to determine that *B. cereus* PelD contains a PelD-like GGDEF domain (Pfam entry PF16963). Only analysis of this sequence by the tertiary structure prediction algorithm implemented in Phyre^2^ [30] was able to link the *B. cereus* PelD sequence to the *P. aeruginosa* PelD structure. This may be linked to the significant architectural variation between Gram-positive PelD orthologs, which suggests a spectrum of activities associated with these proteins that is not limited to c-di-GMP recognition (Fig. S2). In fact, the GGDEF domain is completely absent in Streptococcal PelD orthologs, implying that if Streptococci produce Pel, they must utilize a different mechanism for the post-translational regulation of Pel biosynthesis than that described here for *B. cereus*. Alternatively, it is possible that Pel biosynthesis is not regulated post-translationally in Streptococci. Interestingly, we noted that PelD from *S. thermophilus* JIM8232 contained only the transmembrane region and GAF domain, likely due to interruption of this gene by a transposon (Fig. S2, S6). However, despite this truncated *pelD* gene, JIM8232 produces significant amounts of biofilm biomass that is dependent on the *pel* locus [59]. In contrast, *S. salivarius* JIM8777, which also utilizes the *pel* locus [59], has an intact epimerase region at the amino-terminus of its PelD protein (Fig. S2). This suggests either that the predicted epimerase domain of PelD present in all other Streptococcal strains identified here is not required for biofilm formation in JIM8232, or perhaps in Streptococcal species PelD as a whole is dispensable.

Amongst the Streptococcal, Clostridial, and Actinobacterial species identified in this analysis, the additional epimerase domain at the amino-terminus of PelD correlated with the presence of a well-conserved set of accessory genes located immediately downstream of and contiguous with the *pelDEA_DA_FG* operon (Fig. S4, S5). The protein products of these accessory genes are predicted to encode, among other things, an ortholog of the spore coat protein CotH. While CotH functions as a protein kinase within the spore coat of endospore forming bacteria [35], its identification here amongst non-spore forming Streptococci suggests that, within the context of the putative *pelDEA_DA_FG* locus and the adjoining accessory genes, the *cotH*-like gene may be performing a different function. One possibility is that it may play a role in the regulation of Pel biosynthesis via protein phosphorylation. This hypothesis is particularly relevant in Streptococci as they would not be able to regulate Pel biosynthesis via a canonical c-di-GMP-based mechanism.

Taken together, the data and analyses we have presented here suggests that the Pel polysaccharide is utilized by the divergent Firmicutes *B. cereus, S. thermophilus*, and *S. salivarius* for biofilm formation, and that a diverse array of Gram-positive bacteria have the genetic capacity to produce Pel via a conserved *pel* biosynthesis cluster (Fig. 1). This adds to the already growing list of Gram-negative organisms that have been shown to possess the genetic capacity for Pel biosynthesis [27] and suggests that Pel biosynthesis may be a widespread phenomenon amongst bacteria generally. Based on our analysis of the *pel* operon in *B. cereus* ATCC 10987, we have produced a model for Pel biosynthesis in Gram-positive bacteria (Fig. 7) that exhibits similarity with the current model for Pel production in Gram-negative bacteria (Fig. S1). Furthermore, our analysis of operon architectures in other Gram-positive bacteria suggests that there is still a great deal to be uncovered regarding the mechanism of Pel biosynthesis and its regulation in non-Bacilli. In particular, the functional consequences of the PelD modifications identified here (Fig. S2) and the role, if any, of the downstream accessory genes found in many Gram-positive bacteria (Fig. S4, S5) may reveal details of a c-di-GMP independent mechanism of Pel regulation.

**Figure 7:**
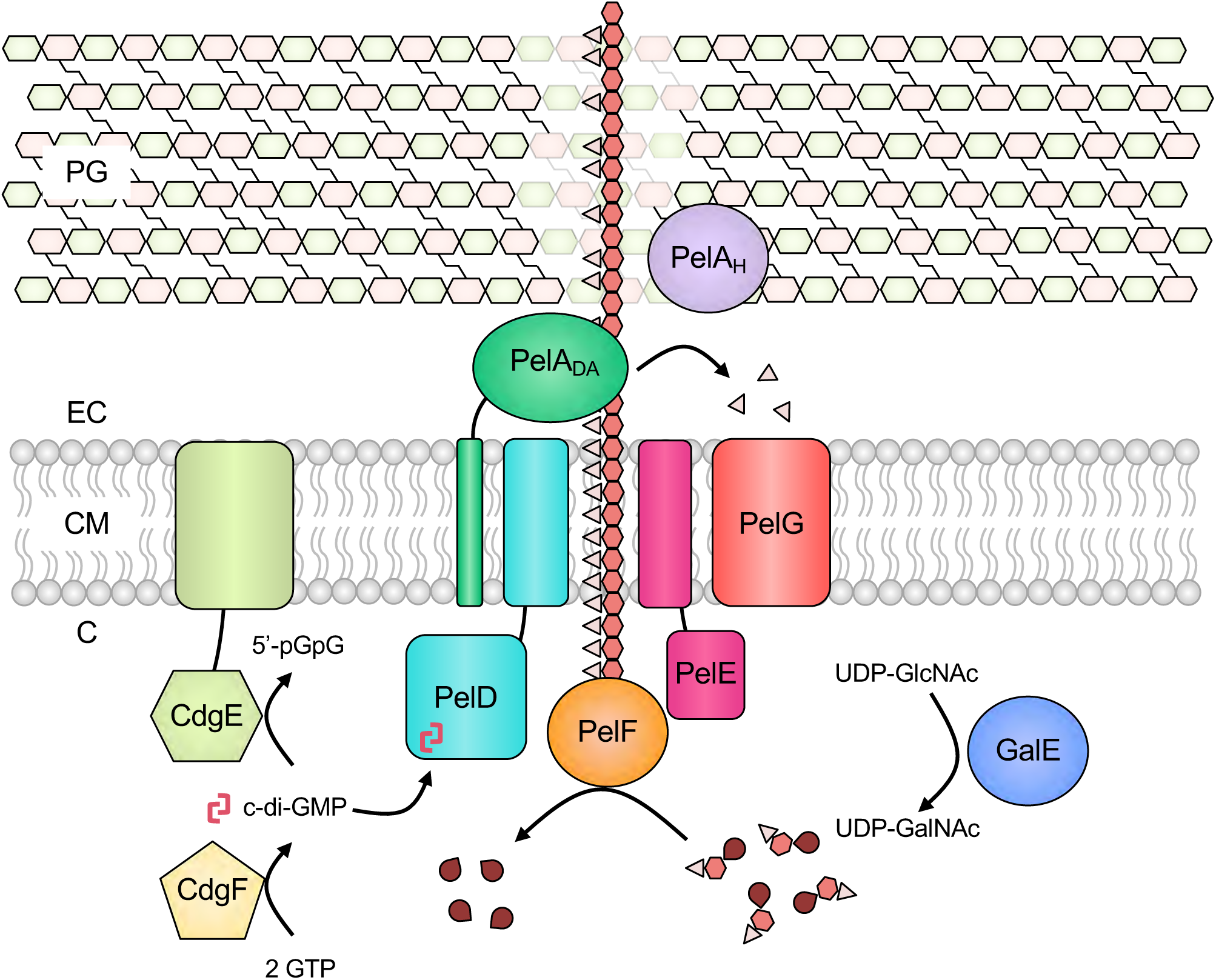
Model for *B. cereus* ATCC 10987 Pel biosynthesis. PelF is a glycosyltransferase that polymerizes UDP-GalNAc to generate the Pel polysaccharide (light pink triangles are acetyl groups, red hexagons are GalN monosaccharide units, dark red teardrops are UDP). Once synthesized, Pel is exported across the cytoplasmic membrane via PelD, PelE, PelA_DA_, and/or PelG. PelA_DA_, corresponding to the deacetylase domain of *P. aeruginosa* PelA, partially deacetylates the polymer to generate mature Pel. PelA_H_, corresponding to the hydrolase domain of *P. aeruginosa* PelA, performs a role in regulating overall biofilm formation levels of ATCC 10987. Pel biosynthesis requires binding of the nucleotide second messenger c-di-GMP to the I-site of the cytoplasmic GGDEF domain of PelD. CdgF is a diguanylate cyclase that synthesizes c-di-GMP from two GTP molecules and thus positively regulates Pel biosynthesis. CdgE is a phosphodiesterase that converts c-di-GMP to 5’-pGpG and thus negatively regulates Pel biosynthesis. GalE epimerizes UDP-GlcNAc to UDP-GalNAc, which is presumably utilized by PelF for Pel polymerization. C, cytoplasm; CM, cytoplasmic membrane; EC, extracellular space; PG, peptidoglycan; UDP, uridine di-phosphate; GlcNAc, *N-*acetylglucosamine; GalNAc, *N*-acetylgalactosamine; GTP, guanosine tri-phosphate; c-di-GMP, cyclic-3’,5’-dimeric guanosine monophosphate; 5’-pGpG, phosphoguanylyl-3’,5’-guanosine.

## Materials and Methods

### Bacterial strains, plasmids and growth conditions

A detailed list of the bacterial strains and plasmids used in this study can be found in Table S2. Generation of plasmids was performed using *E.* coli DH5α as a host for replication. All *B. cereus* mutant strains were derived from ATCC 10987 and were constructed using allelic exchange [61], as described below. Unless otherwise stated, lysogeny broth (LB) was used for growth of all strains. LB contained, per litre of ultrapure water, 10 g tryptone, 5 g NaCl, and 10 g yeast extract. To prepare solid media, 1.5% (w/v) agar were added to LB. Where appropriate, antibiotics were added to growth media. For *E. coli*, 100 μg/mL carbenicillin, 50 μg/mL kanamycin, or 100 μg/mL spectinomycin was used, and for *B. cereus*, 3 μg/mL erythromycin or 10 μg/mL chloramphenicol were used, as necessary.

### Standard molecular methods

All basic microbiological and molecular biological techniques were performed using standard protocols. Genomic DNA was isolated using BioRad InstaGene Matrix. Plasmid isolation and DNA gel extraction was performed using purification kits purchased from BioBasic. Restriction enzymes, DNA ligase, alkaline phosphatase, and DNA polymerase were purchased from Thermo Scientific. Primers used in this study were obtained from Sigma Aldrich (Table S3). Sanger sequencing to confirm the sequence of plasmids and chromosomal mutations (described below) was performed at The Center for Applied Genomics, Toronto.

### Transformation of B. cereus ATCC 10987

To overcome the restriction barrier of *B. cereus* ATCC 10987 and improve transformation efficiency, a plasmid DNA methylation pipeline [62] was employed. Briefly, plasmids to be transferred into *B. cereus* ATCC 10987 were first transformed into *E. coli* EC135, a strain engineered to remove all native DNA methyltransferases [62], alongside the plasmid pM.Bce, which was designed to express DNA methyltransferases present in the ATCC 10987 genome [62]. EC135, containing both pM.Bce and the plasmid of interest to be methylated, were plated on LB agar containing 100 μg/mL spectinomycin and 100 μg/mL carbenicillin and incubated overnight at 30 °C. Transformants were then grown overnight at 30 °C in 5 mL of LB supplemented with 100 μg/mL spectinomycin, 100 μg/mL carbenicillin, and 0.2% (w/v) L-arabinose to induce expression of ATCC 10987 methyltransferases to methylate the plasmid of interest. Methylated plasmids were isolated and transformed into *B. cereus* ATCC 10987, as described below.

To generate electrocompetent *B. cereus*, 2 mL of an overnight LB culture grown to saturation was used to inoculate 500 mL of LB, which was then grown at 37 °C to an OD_600nm_ of 0.2 – 0.4. Cells were collected by centrifugation at 4000 × *g* for 10 min at 4 °C and washed in 10 mL of cold electroporation buffer (1 mM Tris pH 8.0, 10% (w/v) sucrose, 15% (v/v) glycerol). Cells were collected and washed in 10 mL of cold electroporation buffer four more times, as described above. Cells were then resuspended in a final volume of 4 mL of electroporation buffer, divided into 100 μL aliquots, and used immediately or stored at -80 °C for later use.

To transform *B. cereus* ATCC 10987, 0.5 -2 μg of methylated plasmid DNA was mixed with 100 μL of electrocompetent *B. cereus* cells generated as described above, transferred to a pre-chilled 2 mm gap electroporation cuvette (Bio-Rad), and incubated on ice for 10 min. Cells were then pulsed once using a BioRad Gene Pulser electroporator with capacitance extender at 2.5 kV, 200 Ω, and 25 μF, followed by immediate addition of 1 mL of NCMLB medium (100 mM K_2_HPO_4_, 200 mM NaCl, 30 mM glucose, 10 mg/mL tryptone, 5 mg/mL yeast extract, 1 mM trisodium citrate, 0.2 mM MgSO_4_, 380 mM mannitol, 500 mM sorbitol, pH 7.2). Electroporated cells were then transferred to a 1.5 mL tube and grown at 37 °C for 3 h, before being plated onto LB with the appropriate antibiotic.

### Chromosomal mutation in B. cereus ATCC 10987

In-frame, unmarked *pelD*, *pelE*, *pelA_DA_*, *pelF*, *pelG*, *pelA_H_*, *cdgE*, and *cdgF* deletion mutants of *B. cereus* ATCC 10987 were generated using an established allelic exchange protocol [61]. Construction of the gene deletion alleles was performed by amplifying flanking regions of the ORF of the respective gene and joining these flanking regions by splicing-by-overlap extension PCR using Phusion DNA polymerase (Thermo Scientific). The upstream forward and downstream reverse primers (Table S3) were tailed with restriction enzyme cleavage sites to enable ligation-dependent cloning of the spliced PCR products. The assembled Δ*pelD*, Δ*pelE*, Δ*pelA_DA_*, Δ*pelF*, Δ*pelG*, Δ*pelA_H_*, Δ*cdgE*, and Δ*cdgF* alleles were ligated into pMAD [61] using T4 DNA ligase (Thermo Scientific). The resulting allelic exchange vectors (Table S2), were transformed into *E. coli* DH5α and selected on LB agar containing 100 μg/mL carbenicillin. Plasmids were then isolated from individual colonies and verified by Sanger sequencing using pMAD-SEQ-F and pMAD-SEQ-R primers (Table S3).

To generate deletion mutants (Table S2), Δ*pelD*, Δ*pelE*, Δ*pelA_DA_*, Δ*pelF*, Δ*pelG*, Δ*pelA_H_*, Δ*cdgE*, or Δ*cdgF* allelic exchange vectors were transformed into *B. cereus* using the electroporation protocol described above. Electroporated cells were then transferred onto LB agar containing 3 μg/mL erythromycin and 50 μg/mL 5-bromo-4-chloro-3-indolyl-β-D-galactopyranoside (X-Gal) and grown at 30 °C for up to five days, or until blue colonies began to appear corresponding to single crossover mutants [61]. Blue colonies were then pooled and used to inoculate a 5 mL LB culture without antibiotic, which was grown at 37 °C overnight to saturation. Three more cycles of growth were subsequently performed by subculturing the saturated culture into fresh, antibiotic-free LB, and growing again to saturation. Serial dilutions of the final saturated culture were then plated onto LB agar to obtain single colonies, which were then resuspended in 50 μL of sterile water and 2 μL drops were plated onto LB agar with 50 μg/mL of X-Gal and LB agar with 3 μg/mL of erythromycin. For those colonies that appeared white on X-Gal agar and were erythromycin sensitive, colony PCR was performed using primers that targeted the outside, flanking regions of the gene of interest (Table S3) to identify unmarked gene deletions. These PCR products were then Sanger sequenced using the same primers to confirm the correct deletion.

To generate R363A, D366A, and R395A chromosomal point variants (Table S2), the upstream and downstream genomic regions of *pelD* flanking the codon to be mutated were amplified and joined by splicing-by-overlap extension PCR using Phusion DNA polymerase (Thermo Scientific). The upstream reverse and downstream forward primers (Table S3) were tailed with an ∼15-20 bp segment centered on the codon to be mutated and containing the relevant base variations to generate an alanine codon at this position. The upstream forward and downstream reverse primers (Table S3) were tailed with restriction enzyme cleavage sites to enable ligation-dependent cloning of the spliced PCR products as described above for gene deletion alleles. Allelic exchange was subsequently performed as described above, and chromosomal point variants were confirmed by Sanger sequencing of PCR products generated by colony PCR using primers flanking the region of *pelD* that was mutated (Table S3).

### Complementation of B. cereus gene deletions

To express genes in *B. cereus* for complementation of chromosomal deletions, the vector pAD123-P_xyl_ (Table S2) was constructed by amplifying the *xylAR* promoter element from pHCMC04 [63] and cloning this fragment into pAD123 [64] using forward and reverse primers tailed with BamHI and SacI restriction sites, respectively (Table S3). A new multiple cloning site was subsequently constructed immediately downstream of the *xylAR* promoter using primers tailed with restriction enzyme cleavage sites (Table S3) to generate pAD123-P_xyl_.

To generate complementation vectors, the gene of interest was amplified from *B. cereus* ATCC 10987 genomic DNA using Phusion DNA polymerase (Thermo Scientific) and primers tailed with restriction enzyme cleavage sites for ligation-dependent cloning of the amplified gene (Table S3). The forward primer in each case was tailed with the synthetic ribosome binding site 5’-TAAGGAGGAAGCAGGT-3’. The gene of interest was then ligated into pAD123-P_xyl_ using T4 DNA ligase (Thermo Scientific). The resulting complementation vectors (Table S2), were transformed into *E. coli* DH5α and selected on LB agar containing 100 μg/mL carbenicillin. Plasmids were then isolated from individual colonies and verified by Sanger sequencing using pAD123-P_xyl_-SEQ-F and pAD123-P_xyl_-SEQ-R primers (Table S3).

To generate complemented *B. cereus* strains (Table S2), expression vectors were transformed into ATCC 10987 mutants using the protocol outline above, and transformants were selected on LB agar containing 10 μg/mL chloramphenicol. Expression of genes from pAD123-P_xyl_ was induced using 0.001 – 1 % (w/v) xylose, as indicated.

### Crystal violet microtiter plate assay

As an indirect measure of biofilm formation, the crystal violet assay was performed to assess the ability of *B. cereus* mutant strains to adhere to the wells of a plastic 96-well microtiter plate [65]. To make direct comparisons with complemented strains, empty pAD123-P_xyl_ was transformed into wild-type ATCC 10987 and deletion mutants as a vector control, as appropriate, and 10 μg/mL chloramphenicol was added to all cultures. *B. cereus* strains were grown overnight to saturation in LB supplemented with 10 μg/mL chloramphenicol and subsequently normalized to an OD_600nm_ = 0.5. 100 μL of this culture, supplemented with 10 μg/mL chloramphenicol and 0.001 – 1 % xylose, was added to the wells of a Corning CellBind 96-well microtiter plate in triplicate and incubated for 24 h at 30 °C in a humidified chamber to allow cells to adhere to the walls of the plate. Bacterial culture was then gently removed and each well was washed with 200 μL of water twice, followed by the addition of 125 μL of 0.1% (w/v) crystal violet and incubation at room temperature for 10 min. The crystal violet solution was removed and each well was washed with 200 μL of water three times before the plate was allowed to air dry for at least 2 hours. 125 μL of 30% (v/v) acetic acid was subsequently added to each well of the plate and incubated at room temperature for 10 min to solubilize any crystal violet bound to adherent biomass. 100 μL of solubilized crystal violet solution from each well was then transferred to a clean, standard 96-well microtiter plate (Grenier) and absorbance was measured at 550 nm.

### Pellicle production assay

To assess the ability of *B. cereus* strains to form an air-liquid interface biofilm, or pellicle, 5 mL of LB was inoculated with *B. cereus* and incubated overnight at 37 °C with shaking. To make direct comparisons with complemented strains, empty pAD123-P_xyl_ was transformed into wild-type ATCC 10987 and deletion mutants as a vector control, as appropriate, and 10 μg/mL chloramphenicol was added to all cultures. The next day, the density of the culture was adjusted to OD_600nm_ = 1.0 and 30 μL of the resulting culture was used to inoculate 3 mL of LB containing 0.001 - 1% (w/v) xylose and 10 μg/mL of chloramphenicol in borosilicate glass tubes. The glass tubes were left undisturbed at 30 °C for 24 h, at which point the air-liquid interface was photographed. Pellicle formation was indicated by complete coverage of the air-liquid interface by an opaque layer of cells.

To visualize biomass adherent to the walls of borosilicate glass tubes, pellicles were generated as described above and media was gently removed by pipetting. The glass tube was washed twice with 4 mL of water, followed by staining for 10 min with 4 mL of 0.1% (w/v) crystal violet at room temperature. The crystal violet was then gently removed, followed by three washes with 4 mL water. The adherent biomass stained by crystal violet was then photographed.

### Scanning electron microscopy

For SEM, overnight cultures of *B. cereus* ATCC 10987 (wild-type and Δ*pelF*) were diluted to 10^5^ cfu/mL. 96-well plates were inoculated with 150 μL of this diluted culture and biofilms were grown on suspended plastic coverslips for 24 h at 37 °C. Biofilms were formed at the air-liquid interface. The preparation for SEM was done as reported previously and is briefly described here [66]. Coverslips carrying *B. cereus* biofilms were washed with a pre-fixation solution (0.075% Ruthenium Red, 2.5% glutaraldehyde, 50 mM lysine in 100 mM HEPES pH 7.3) for 30 min followed by fixation for 1 h (0.075% Ruthenium Red, 2.5% glutaraldehyde in 100 mM HEPES pH 7.3). Coverslips were washed briefly in HEPES (100 mM, pH 7.4), and then incubated with 2% osmium tetroxide (OsO_4_) for 30 min. A quick dehydration series was performed in ethanol at 50, 70, 80, 90 and 100% for 10 min each, followed by 3 changes of 100% ethanol. Samples were then critical point dried by replacing the ethanol with CO_2_ using the Denton DCP-1 and sputter coated in gold using the Denton Desk V TSC. Samples then imaged using the Quanta FEG 250 SEM, operated at 10 kV under high vacuum.

### Epifluorescence microscopy

For fluorescence microscopy, cultures were diluted from overnight cultures to 10^5^ cfu/mL and inoculated into 96-well plates in the same manner as for SEM biofilm growth. Glass coverslips were coated in 0.01% poly-ι-lysine and placed in the inoculated wells where biofilms were allowed to form for 24 h at 37 °C. 24-h-old biofilms were washed twice in PBS) and then sequentially incubated with 100 μg/mL DAPI and 100 μg/mL WFL for 15 min each. Biofilms were washed briefly 3 times in PBS before being mounted on glass slides. Labelled biofilms were imaged using an inverted Axiovert 200M fluorescence microscope (Zeiss) equipped with an ORCA-R^2^ C10600 digital camera (Hamamatsu).

### Dot blots

Pel antisera was obtained as described in Colvin *et al* from *P. aeruginosa* PA14 P_BAD_*pel* [31]. The adsorption reaction was conducted as described by Jennings *et al* [41]. Cell associated Pel was obtained as previously described in Colvin *et al* [31]. Cells were harvested by centrifugation (16,000 × *g* for 2 min) from 1 mL aliquots of *B. cereus* ATCC 10987 grown overnight at 37 °C in LB. The supernatant containing secreted Pel was set aside and cell pellets were resuspended in 100 μl of 0.5 M EDTA, pH 8.0. Cells were boiled for 20 min with occasional vortexing, and centrifuged (16,000 × *g* for 10 min) to harvest the supernatant containing cell associated Pel. Cell-associated and secreted Pel were treated with proteinase K (final concentration, 0.5 mg/mL) for 60 min at 60 °C, followed by 30 min at 80 °C to inactivate proteinase K.

Pel immunoblots were performed as described by Colvin *et al* [31] and Jennings *et al* [41]. 5 μL of cell associated and secreted Pel, prepared as described above, were pipetted onto a nitrocellulose membrane and left to air dry for 10 min. The membrane was blocked with 5% (w/v) skim milk in Tris-buffered saline with Tween-20 (TBS-T) for 1 h at room temperature and probed with adsorbed α-Pel at a 1:60 dilution in 1% (w/v) skim milk in TBS-T overnight at 4 °C with shaking. Blots were washed three times for 5 min each with TBS-T, probed with goat α-rabbit HRP-conjugated secondary antibody (Bio-Rad) at a 1:2000 dilution in TBS-T for 45 min at room temperature with shaking, and washed again. All immunoblots were developed using SuperSignal West Pico (Thermo Scientific) following the manufacturer’s recommendations.

For WFL-HRP immunoblots, 5 μL of cell associated and secreted Pel, prepared as described above, were pipetted onto a nitrocellulose membrane and left to air dry for 10 min. The membrane was blocked with 5% (w/v) bovine serum albumin (BSA) in TBS-T for 1 h at room temperature and probed with 10 μg/mL of WFL-HRP (EY Laboratories) in 2% (w/v) BSA in TBS-T with 0.2 g/L CaCl_2_ overnight at room temperature with shaking. Membranes were washed twice for 5 min and once for 10 min with TBS-T, then developed as described above.

### Cloning, expression, and purification of PelA_H_^Bc^ and PelA_H_<u>^Pa^</u>

To generate a plasmid for expression of PelA_H_^Bc^ (BCE_5582) in *E. coli*, the *pelA_H_^Bc^* ORF, excluding the predicted signal sequence (residues 1-21), was amplified from *B. cereus* ATCC 10987 genomic DNA with Phusion DNA polymerase (Thermo Scientific) using primers tailed with restriction enzyme cleavage sites for ligation-dependent cloning (Table S3). The gene of interest was then ligated into pET24a (Novagen), which encodes a C-terminal hexahistidine-tag, using T4 DNA ligase (Thermo Scientific). The resulting PelA_H_^Bc^ expression vector (Table S2), was transformed into *E. coli* DH5α and selected on LB agar containing 50 μg/mL kanamycin. Plasmids were then isolated from individual colonies and verified by Sanger sequencing using T7 and T7ter primers (Table S3).

To generate the E213A point variant of PelA_H_^Bc^, the upstream and downstream regions of *pelA_H_^Bc^* flanking E213 were amplified and joined by splicing-by-overlap extension PCR using Phusion DNA polymerase (Thermo Scientific). The upstream reverse and downstream forward primers (Table S3) were tailed with an ∼15-20 bp segment centered on E213 that contained the relevant base variations to generate an alanine codon at position 213. The upstream forward and downstream reverse primers (Table S3) were tailed with restriction enzyme cleavage sites to enable ligation-dependent cloning of the spliced PCR product as described above for wild-type PelA_H_^Bc^.

For expression of PelA_H_^Bc^, pET24a::PelA_H_^Bc^ was transformed into *E. coli* BL21 CodonPlus^TM^ (DE3) competent cells (Stratagene). Cells were grown at 37 °C until the OD_600nm_ of the culture reached 0.5, at which point the temperature was shifted to 18 °C and 1 mM of isopropyl β-D-1-thiogalactopyranoside (IPTG) was added to induce expression of PelA_H_^Bc^. Cells were grown overnight at 18 °C and harvested the next day via centrifugation at 5,000 × *g* for 20 min at 4 °C. Cell pellets were used immediately for purification.

To purify PelA_H_^Bc^, cell pellets were resuspended in 50 mL of Buffer A (50 mM Tris-HCl pH 8.0, 500 mM NaCl, 10% (v/v) glycerol), containing one SIGMAFAST EDTA-free protease inhibitor tablet, per 1 L of original culture. Resuspended cells were then lysed by three passes through an Emulsiflex-C3 (Avestin Inc.) at a pressure of 10,000 - 15,000 psi. Insoluble material from the cell lysate was removed via centrifugation at 30,000 × *g* for 30 min at 4 °C. The clarified lysate was then applied to a 5 mL bed Ni^2+^-NTA column that had been pre-equilibrated with ten column volumes of Buffer A containing 10 mM imidazole. Non-specifically bound material was washed from the column using six column volumes of Buffer A containing 20 mM imidazole. Bound protein was subsequently eluted using 3 column volumes of Buffer A containing 250 mM imidazole. The Ni^2+^-NTA-purified protein was then concentrated to 2 mL using an Amicon Ultra centrifugal filtration device (Millipore) with a 10 kDa molecular weight cut-off. PelA_H_^Bc^ was further purified and buffer-exchanged into Buffer B (20 mM Tris–HCl pH 8.0, 150 mM NaCl, 5% (v/v) glycerol) by size-exclusion chromatography using a HiLoad 16/60 Superdex 75 prep-grade gel filtration column (GE Healthcare). PelA_H_^Bc^ eluted as a single Gaussian-shaped peak, and all PelA_H_^Bc^ -containing fractions were pooled and concentrated to 7.5 mg/mL using an Amicon Ultra centrifugal filtration device (Millipore) with a 10 kDa molecular weight cut-off. Concentrated PelA_H_^Bc^ could then be stored at 4 °C for up to two weeks without losing activity. PelA_H_^Bc^-E213A was purified as described above for PelA_H_^Bc^.

Cloning, expression, and purification of PelA_H_*^Pa^*and the E218A mutant has been described previously [44].

### Cloning, expression, and purification of recombinant B. cereus PelD

The nucleotide sequence of *pelD* from *B. cereus* ATCC 10987 (*BCE_5583; WP_000559343.1*) was obtained from the National Center for Biotechnology Information (NCBI) genome database and used to design primers specific to *BCE_5583*. The PCR product of *BCE_5583* was amplified from genomic DNA, digested with NheI and XhoI, and cloned into a pET-28a vector (Novagen). The resulting expression vector pET28a::PelD_150-407_ encodes residues 150-407 of PelD^Bc^ fused to a cleavable N-terminal His_6_-tag (NHis_6_) for purification purposes. The PelD^Bc^ mutants R363A, D366A, and R395A were generated using the QuikChange Lightning site-directed mutagenesis kit (Stratagene). The sequence of all vectors was confirmed using DNA sequencing (The Center for Applied Genomics).

Expression and purification of PelD^Bc^ was performed as described previously for *P. aeruginosa* PelD, with some modifications as outlined below [67]. Expression of PelD^Bc^ was achieved through transformation of pET28a::PelD_150-407_ into *E. coli* BL21 CodonPlus^TM^ (DE3) competent cells (Stratagene), which was then grown in 1 L LB containing 50 µg/mL kanamycin at 37 °C. The cells were grown to an OD_600nm_ of 0.6, whereupon IPTG was added to a final concentration of 1.0 mM to induce expression. The induced cells were incubated for 20 h at 18 °C prior to being harvested via centrifugation at 6,260 × *g* for 20 min at 4 °C. The resulting cell pellet was stored at -20 °C until required.

To purify NHis_6_-PelD^Bc^, and the mutant proteins, the cell pellet from 1 L of bacterial culture was thawed and resuspended in 40 mL of buffer C (50 mM HEPES-HCl pH 8.0, 300 mM NaCl, 5% (v/v) glycerol, 1 mM tris(2-carboxylethyl)phosphine hydrochloride (TCEP)) with 1 SIGMAFAST protease inhibitor EDTA-free cocktail tablet (Sigma). Cells were lysed and NHis_6_-PelD^Bc^ was purified by Ni^2+^-NTA chromatography as described previously for *P. aeruginosa* PelD [67]. SDS-PAGE analysis revealed that the resulting NHis_6_-PelD^Bc^ was 90% pure and appeared at the expected molecular weight of 32.7 kDa. Fractions containing protein were pooled and concentrated to a volume of 2 mL by centrifugation at 2,200 × *g* at a temperature of 4 °C using an Amicon Ultra centrifugal filter device (Millipore) with a 10 kDa molecular weight cut-off (MWCO). The protein was buffer exchanged into buffer D (20 mM HEPES-HCl pH 8.0, 150 mM NaCl, 5% (v/v) glycerol, 1 mM dithiolthreitol (DTT)) by size-exclusion chromatography using a HiLoad 16/60 Superdex 200 gel-filtration column (GE Healthcare). All PelD proteins eluted as single Gaussian peaks. Protein containing fractions were pooled and the protein concentrated by centrifugation at 2,200 × *g* at 4 °C using an Amicon Ultra centrifugal filter device (Millipore) with a 10 kDa MWCO.

### Biofilm disruption assays

Biofilm disruption assays using glycoside hydrolases have been described previously [44, 45]. Briefly, biofilm cultures were established in 96-well microtiter plates as described above. Following incubation, nonadherent cells and media were removed by washing the plate twice with 200 μL of water. The wells were filled with 142.5 μL of PBS followed by 7.5 μL of varying concentrations of PelA_H_*^Bc^*, PelA_H_*^Pa^*, or their respective catalytically inactive point variants (0.4 to 60 μM). Reactions were incubated for 60 min at room temperature on a rotating nutator, at which time the reaction was quenched by washing the plates two times with 200 μL of water. The wells were stained with 125 μL of 0.1% (w/v) crystal violet for 10 min, followed by three washes with water and solubilization of the crystal violet with 30% (v/v) acetic acid. 100 μL of solubilized crystal violet was transferred to a clean 96-well plate and absorbance was quantified at 550 nm. All reactions were completed in triplicate.

### Preparation of c-di-GMP

The protocol for the enzymatic preparation of c-di-GMP was adapted from Zahringer *et al* [68] and De *et al* [69], and performed as described previously [26].

### Isothermal titration calorimetry

For ITC experiments, PelD protein samples and c-di-GMP were prepared in 20 mM HEPES pH 8.0, 150 mM NaCl, 5% glycerol, and 1 mM DTT. ITC measurements were performed with a MicroCal Auto-iTC200 microcalorimeter (Malvern Instruments, Inc. Northampton, MA). Titrations were carried out with 450 µM c-di-GMP in the syringe and a 25 µM solution of the indicated PelD sample in the cuvette. Each titration experiment consisted of thirty-eight 1 µL injections with 180 s intervals between each injection. The heats of dilution for titrating c-di-GMP into buffer were subtracted from the sample data prior to analysis. The ITC data were analyzed using the Origin v7.0 software (MicroCal Inc.) and fit using a single-site binding model.

## Funding Information

This work was supported in part by grants from the Canadian Institutes of Health Research (CIHR; MOP 43998 and FDN154327 to PLH, PJT156111 to CMK) and the Natural Sciences and Engineering Research Council (RGPIN-2014-06664 and RGPIN-2019-06852 to JP). PLH is a recipient of a Canada Research Chair. GBW and LSM have been supported by graduate scholarships from the Natural Sciences and Engineering Council of Canada (NSERC). GBW has been supported by a graduate scholarship from Cystic Fibrosis Canada. LSM has been supported by graduate scholarships from the Ontario Graduate Scholarship Program, and The Hospital for Sick Children Foundation Student Scholarship Program.

---

**Figure S1:**
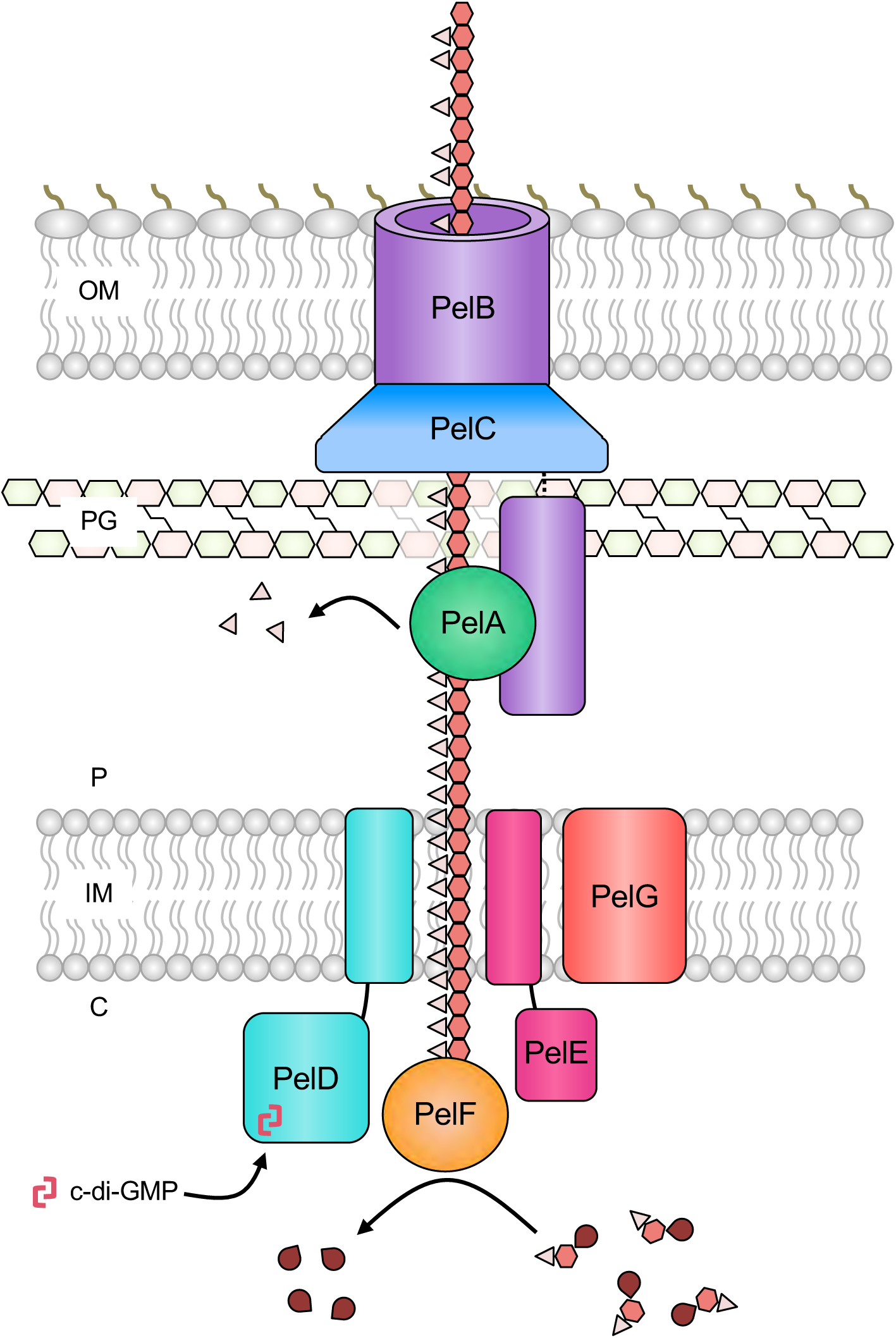
Model of *P. aeruginosa* Pel biosynthesis. PelF is a glycosyltransferase that polymerizes UDP-GalNAc and UDP-GlcNAc to generate the Pel polysaccharide (light pink triangles are acetyl groups, red hexagons are GalN/GlcN monosaccharide units, dark red teardrops are UDP). Once synthesized, Pel is transported across the inner membrane via PelD, PelE, and/or PelG. In the periplasm, PelA interacts with PelB to partially deacetylate the polymer and generate mature Pel, which is subsequently exported across the outer membrane through the β-barrel porin domain of PelB. PelC forms a dodecameric funnel that helps to channel Pel towards the PelB pore. Pel biosynthesis requires binding of the nucleotide second messenger c-di-GMP to the inhibitory site of the cytoplasmic GGDEF domain of PelD. C, cytoplasm; IM, inner membrane; P, periplasmic space; PG, peptidoglycan; OM, outer membrane; c-di-GMP, cyclic-3’,5’-dimeric guanosine monophosphate.

**Figure S2:**
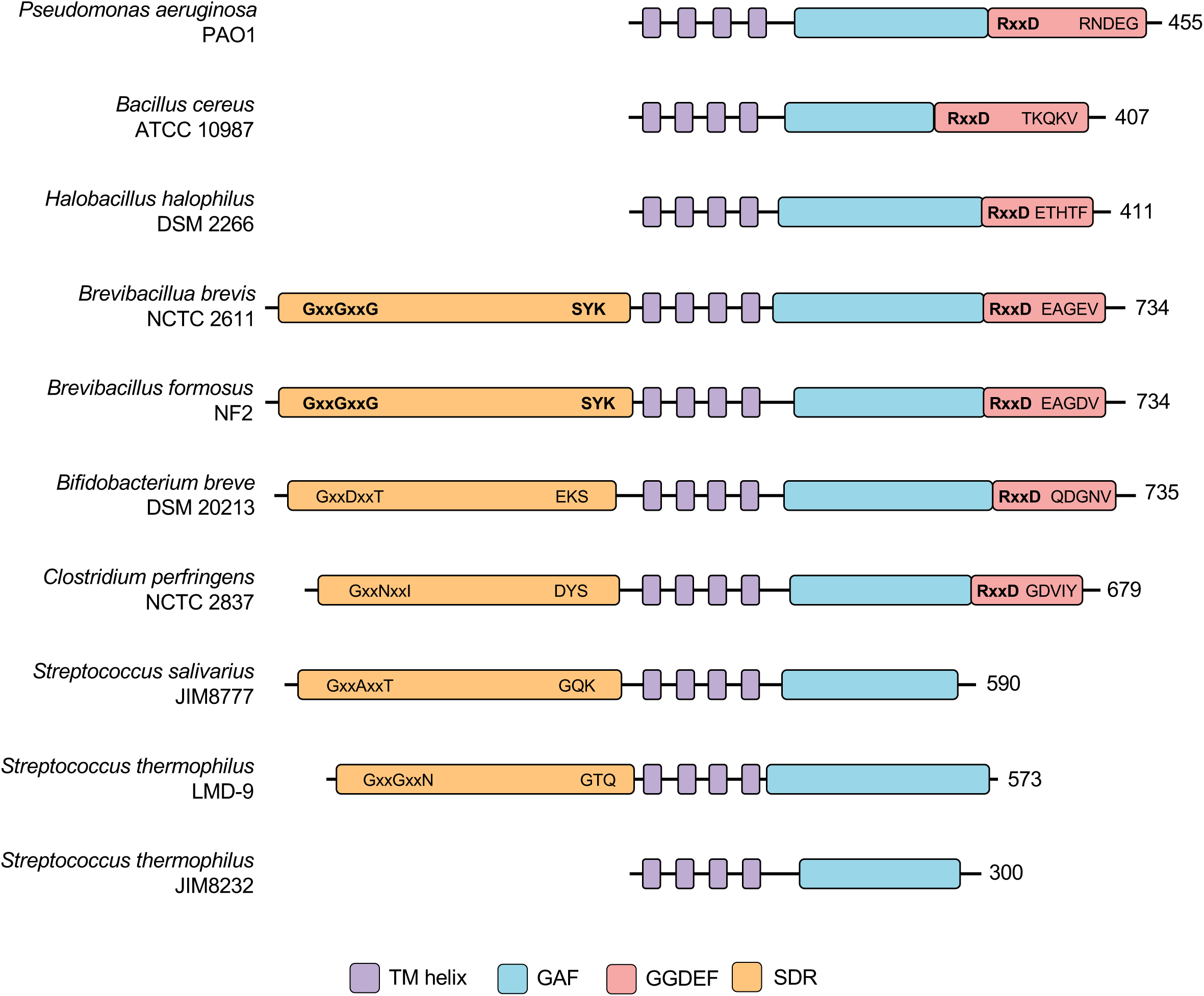
Predicted domain architectures of representative PelD proteins. Predicted domains are depicted as rectangles. Individual domains and overall domain architectures are drawn to scale. The length of each protein is indicated to the right of the respective diagram. Rectangle colours correspond to predicted domain functions, which are listed in the legend at the bottom. Residues from SDR fold that are required for co-factor binding (left) and epimerase activity (right) are shown within the orange rectangle, and are highlighted in bold if they match the consensus sequence for the prescribed function. Residues from the GGDEF fold that are required for c-di-GMP binding to the inhibitory site (left) and diguanylate cyclase activity (righ) are shown within the red rectangle, and are highlighted in bold if they match the consensus sequence required for the prescribed function. TM, transmembrane; GAF, cGMP-specific phosphodiesterase, adenylyl cyclase and FhlA domain; GGDEF, diguanylate cyclase-like domain; SDR, short chain dehydrogenase/reductase.

**Figure S3:**
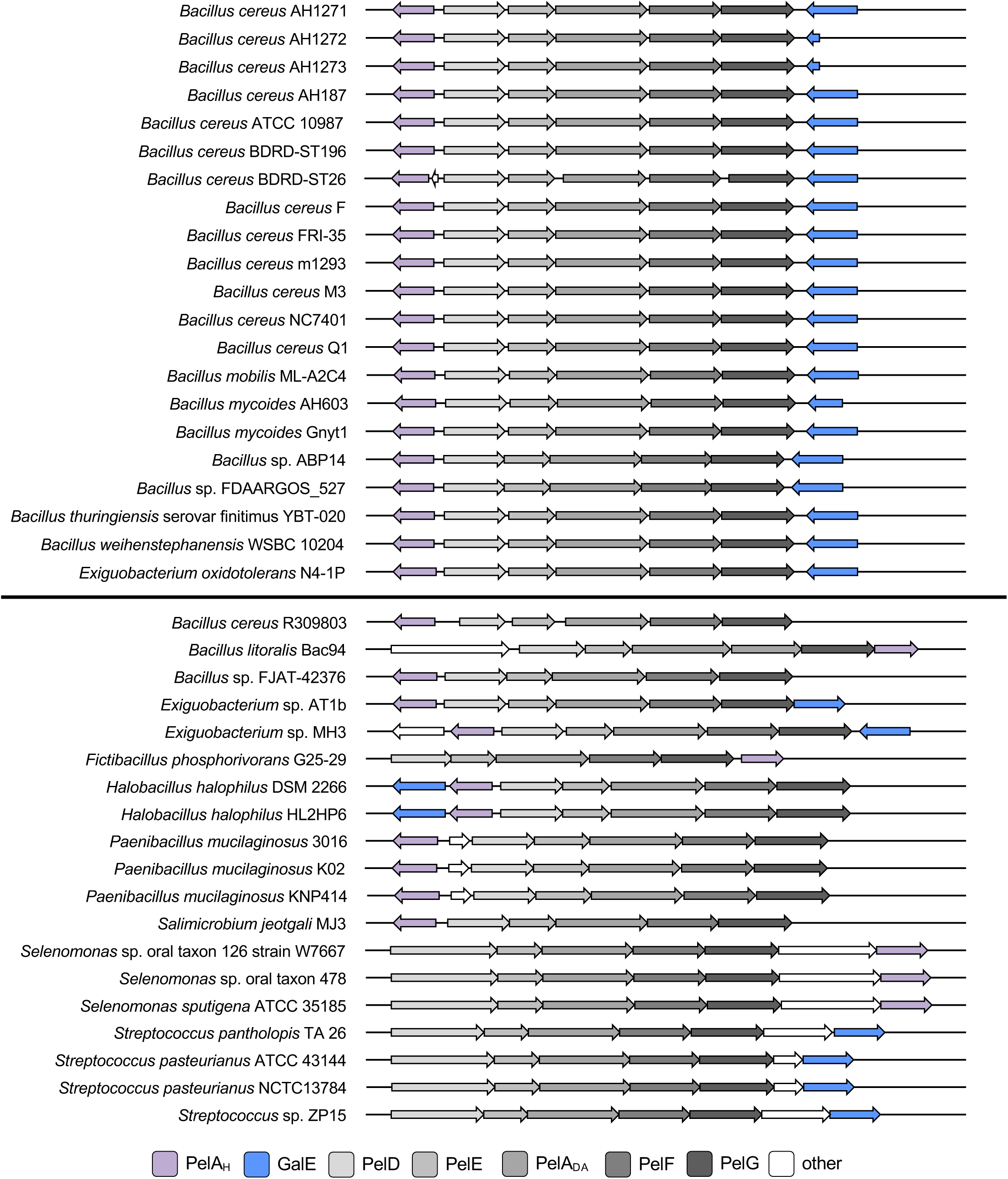
Representative operon architectures containing the accessory genes *galE* and *pelA_H_*. Operons with a conserved accessory gene architecture identified in Bacilli are shown at the top, while operons with divergent architectures are shown at the bottom. Open reading frames are represented as arrows, with the directionality of transcription indicated by the arrow direction. Open reading frames and overall operon architectures are drawn to scale. Arrow colours correspond to predicted protein functions, which are listed in the legend at the bottom. PelA_H_, PelA hydrolase-like domain; PelA_DA_, PelA deacetylase-like domain.

**Figure S4:**
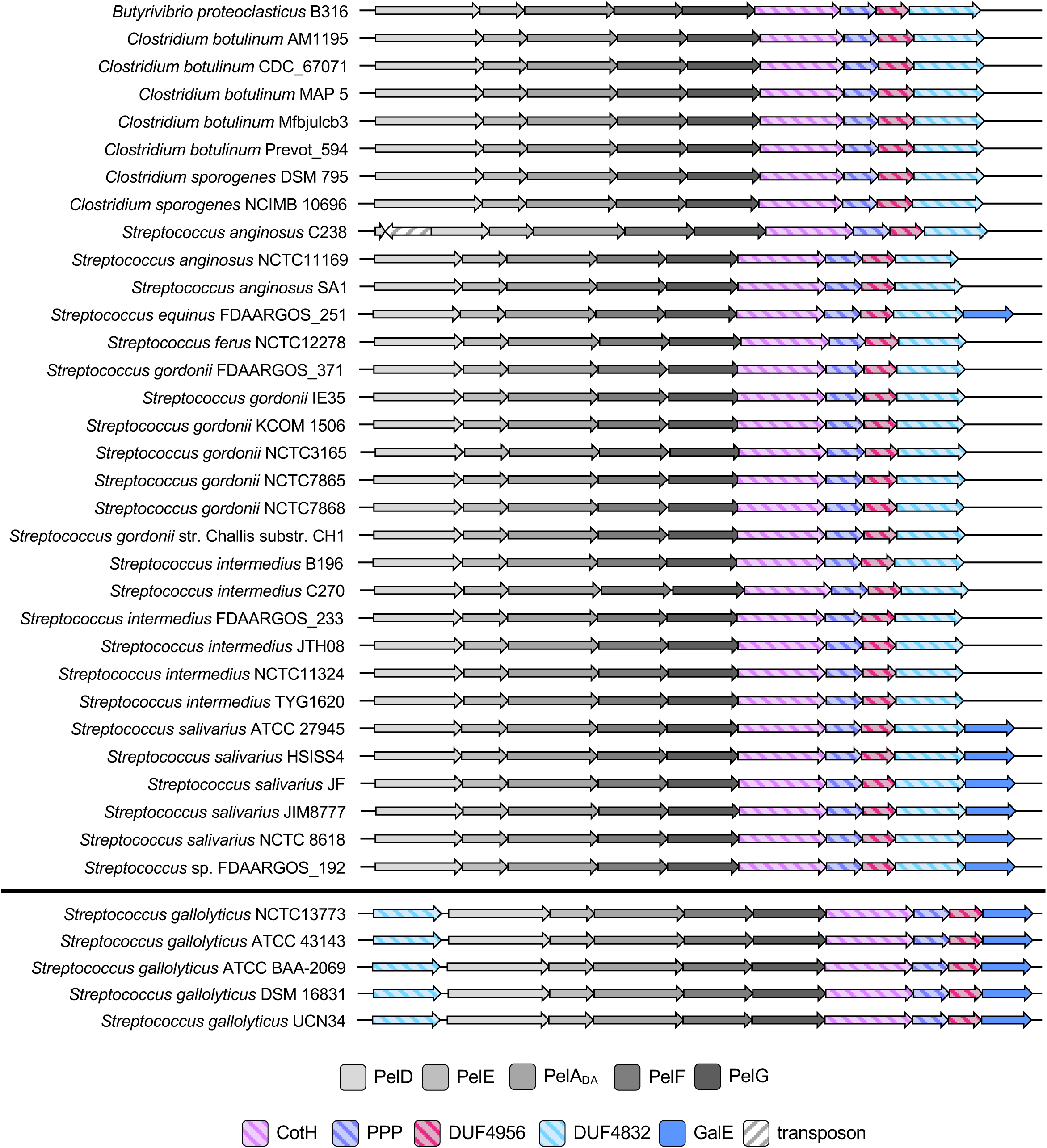
Representative operon architectures identified in Streptococci and Clostridia containing the accessory genes *cotH*, *PPP, DUF4956*, and *DUF4832*. Operons with a conserved accessory gene architecture are shown at the top, while operons with divergent architectures are shown at the bottom. Open reading frames are represented as arrows, with the directionality of transcription indicated by the arrow direction. Open reading frames and overall operon architectures are drawn to scale. Arrow colours correspond to predicted protein functions, which are listed in the legend at the bottom. PelA_DA_, PelA deacetylase-like domain; PPP, poly-phosphate polymerase; DUF, domain of unknown function.

**Figure S5:**
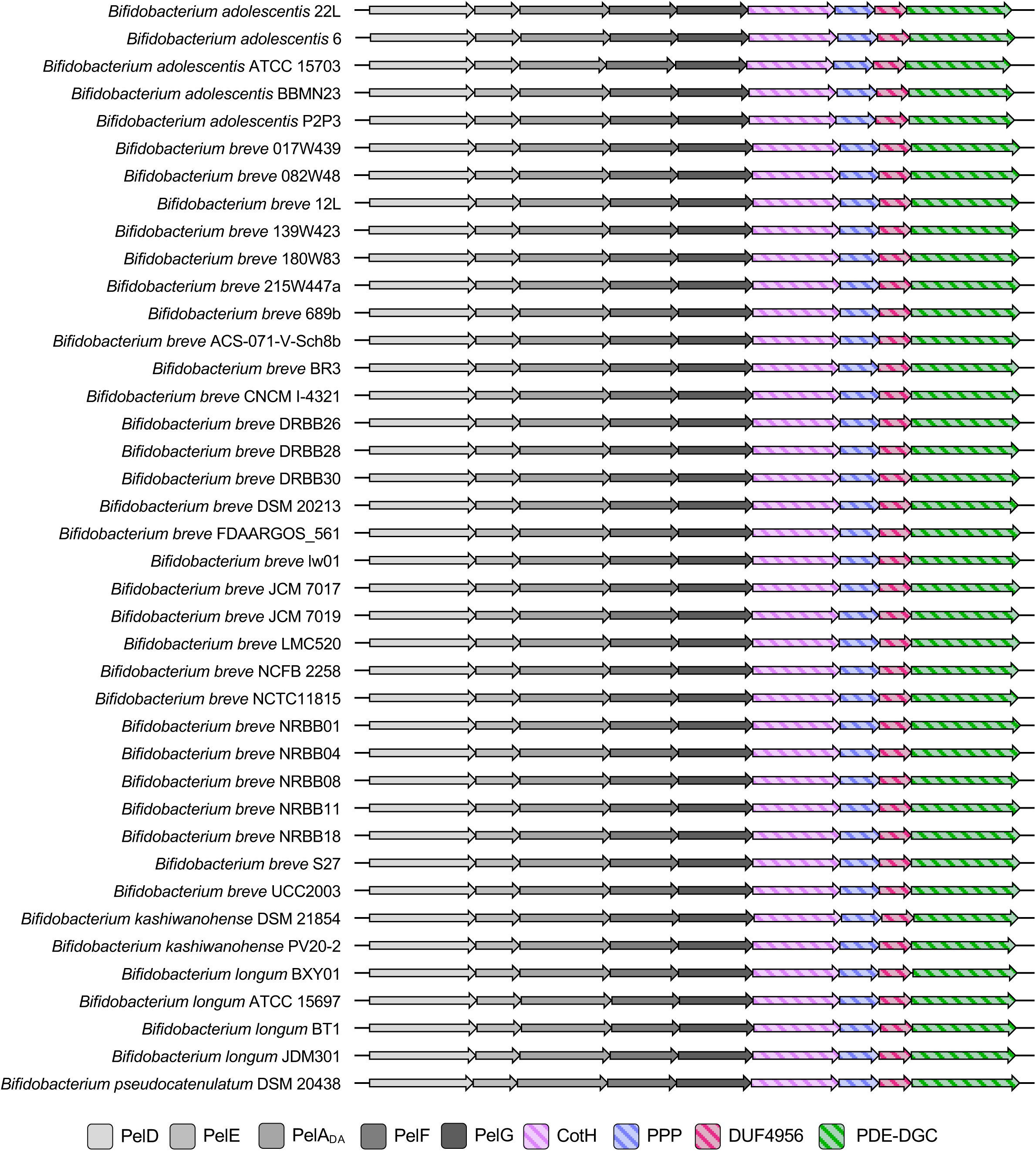
Representative operons with a conserved architecture identified in Bifidobacteria containing the accessory genes *cotH*, *PPP, DUF4956*, and *DGC/PDE*. Open reading frames are represented as arrows, with the directionality of transcription indicated by the arrow direction. Open reading frames and overall operon architectures are drawn to scale. Arrow colours correspond to predicted protein functions, which are listed in the legend at the bottom. PelA_DA_, PelA deacetylase-like domain; PPP, poly-phosphate polymerase; DUF, domain of unknown function; PDE-DGC, phosphodiesterase/diguanylate cyclase-like domain.

**Figure S6:**
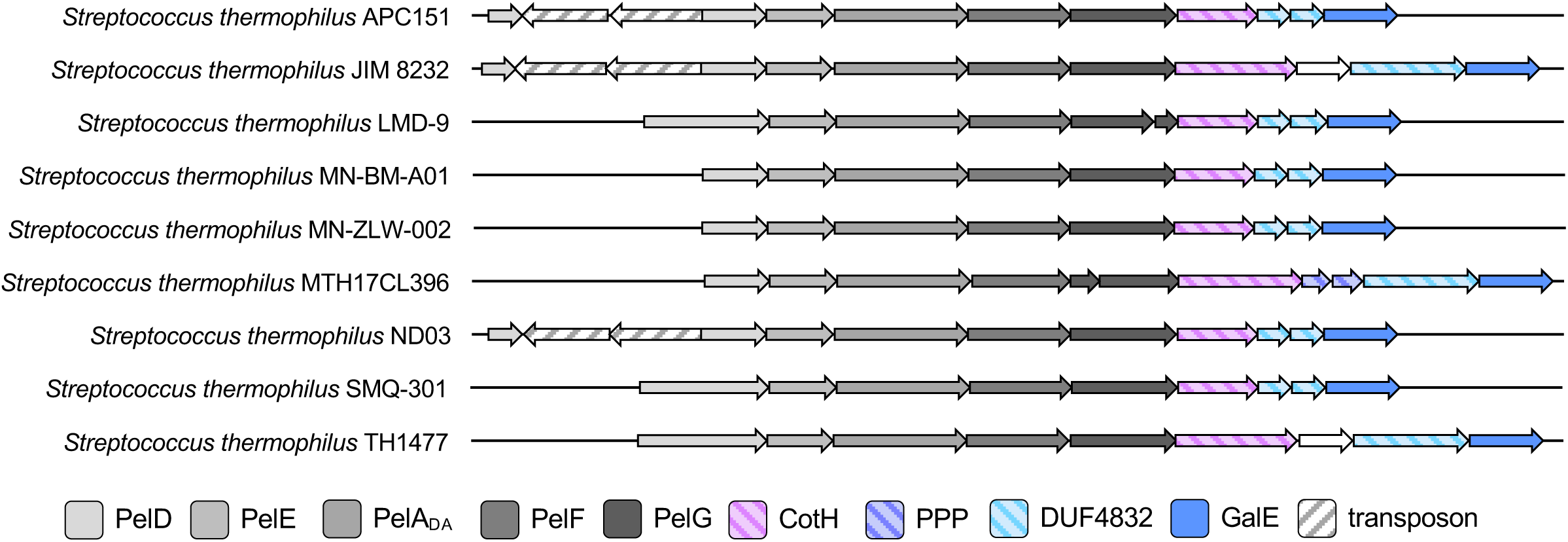
Operon architectures from *Streptococcus thermophilus* strains demonstrating the divergent architectures identified in this species. Open reading frames are represented as arrows, with the directionality of transcription indicated by the arrow direction. Open reading frames and overall operon architectures are drawn to scale. Arrow colours correspond to predicted protein functions, which are listed in the legend at the bottom. PelA_DA_, PelA deacetylase-like domain; PPP, poly-phosphate polymerase; DUF, domain of unknown function.

**Figure S7:**
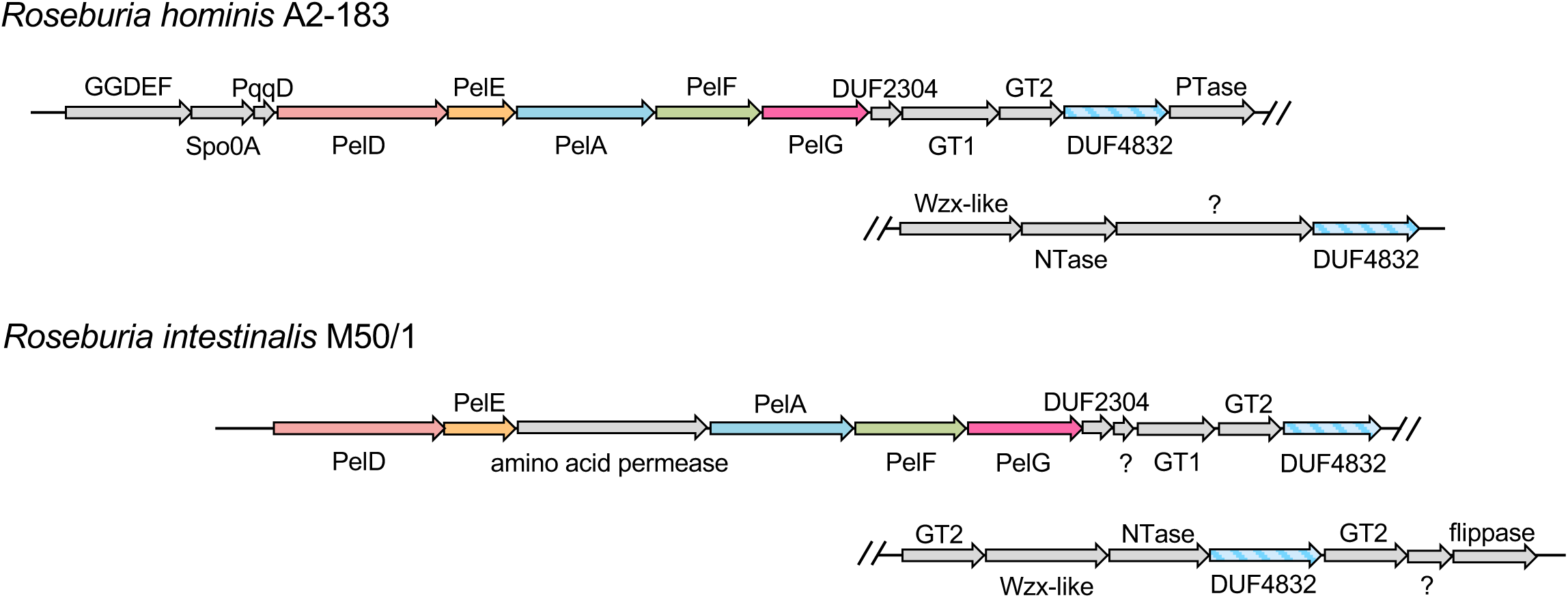
Operon architectures from *Roseburia* species which have the putative *pel* genes within the context of a larger operon likely involved in capsule or teichoic acid biosynthesis. Open reading frames are represented as arrows, with the directionality of transcription indicated by the arrow direction. Open reading frames and overall operon architectures are drawn to scale. The predicted functions of the protein products of each gene are indicated above or below that gene. GGDEF, diguanylate cyclase-like domain; DUF, domain of unknown function; GT1, glycosyltransferase family 1; GT2, glycosyltransferase family 2; PTase, phosphotransferase; NTase, nucleotidyltransferase.

**Figure S8:**
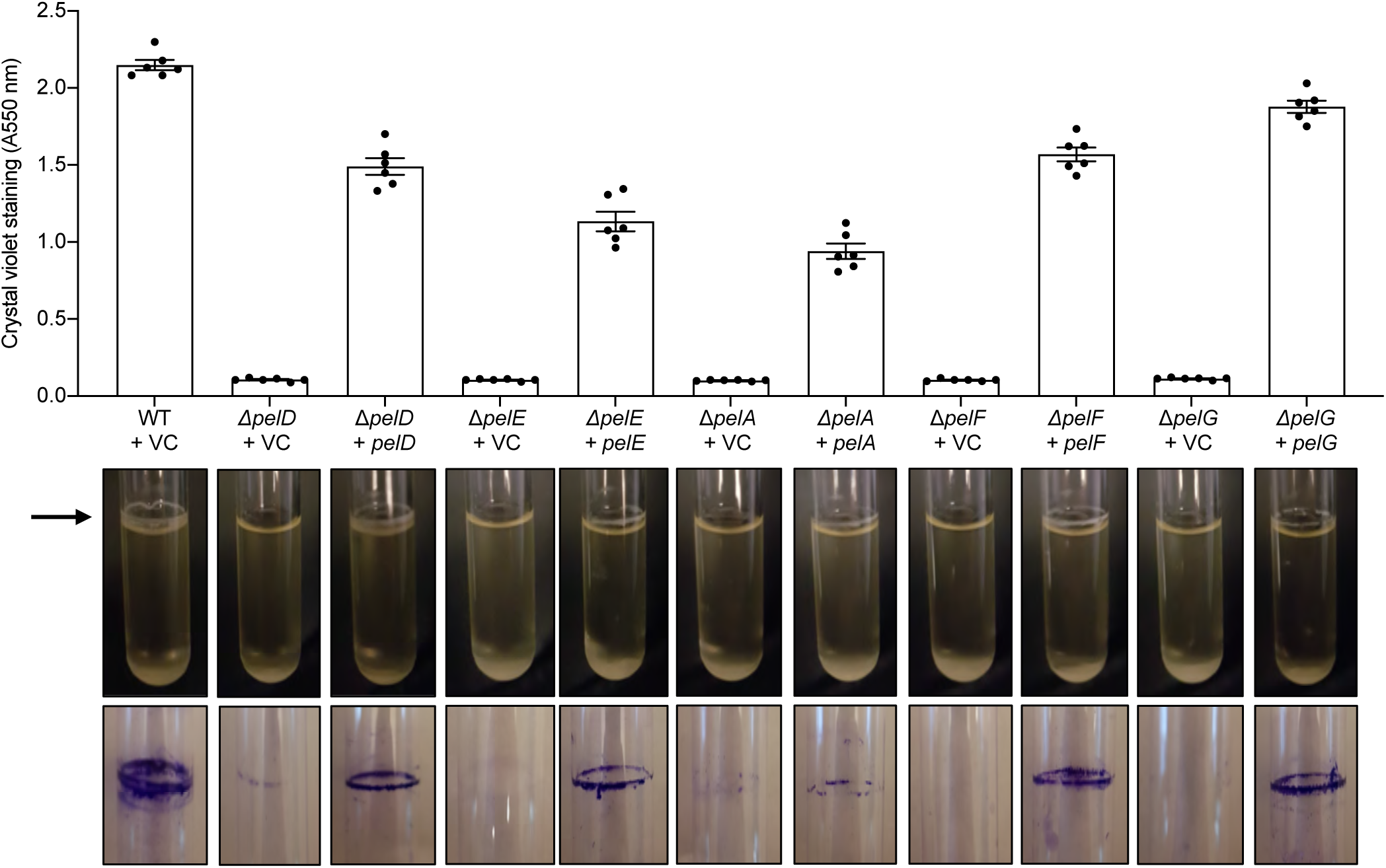
Complementation of *pelDEA_DA_FG* deletion mutants restores biofilm formation. (*top*) Biofilm formation by the indicated strains of *B. cereus* ATCC 10987 assessed by the crystal violet assay. Error bars represent the standard error of the mean of six independent trials. VC, empty vector control. (*middle*) Air-liquid interface (pellicle) biofilm formation by the indicated strains of *B. cereus* ATCC 10987 in borosilicate glass tubes. The pellicle is indicated by the black arrow. (*bottom*) Staining of biomass adherent to the walls of the borosilicate glass tubes pictured above with crystal violet. Non-adherent cells and media were washed from the glass tube before crystal violet staining.

