## Supplementary material for "Discovery and characterization of a Gram-positive Pel polysaccharide biosynthetic gene cluster": SI Tables

**Table S1: Complete list of Gram-positive organisms with *pel*-like operons.**

| **Species*** | **PelG Protein Accession** | **Genome Accession** |
| --- | --- | --- |
| *Bacillus cereus* AH1271 | WP_000907287.1 | NZ_CM000739.1 |
| *Bacillus cereus* AH1272 | WP_000907285.1 | NZ_CM000740.1 |
| *Bacillus cereus* AH1273 | WP_000907285.1 | NZ_CM000741.1 |
| *Bacillus cereus* AH187 | WP_000907295.1 | NC_011658.1 |
| ***Bacillus cereus* ATCC 10987** | **WP_000907294.1** | **NC_003909.8** |
| *Bacillus cereus* BDRD-ST196 | WP_033709612.1 | NZ_CM000725.1 |
| *Bacillus cereus* BDRD-ST26 | EEK97794.1 | CM000724.1 |
| *Bacillus cereus* F | WP_016719067.1 | NZ_CM001787.1 |
| *Bacillus cereus* FRI-35 | WP_014893388.1 | NC_018491.1 |
| *Bacillus cereus* m1293 | WP_000907291.1 | NZ_CM000714.1 |
| *Bacillus cereus* M3 | WP_016719067.1 | NZ_CP016316.1 |
| *Bacillus cereus* NC7401 | WP_000907295.1 | NC_016771.1 |
| *Bacillus cereus* Q1 | WP_000907290.1 | NC_011969.1 |
| *Bacillus cereus* R309803 | EEK76039.1 | CM000720.1 |
| ***Bacillus litoralis* Bac94** | **WP_121663553.1** | **NZ_CP033043.1** |
| ***Bacillus mobilis* ML-A2C4** | **WP_000907287.1** | **NZ_CP031443.1** |
| ***Bacillus mycoides* AH603** | **WP_033707410.1** | **NZ_CM000737.1** |
| *Bacillus mycoides* Gnyt1 | WP_002167287.1 | NZ_CP020743.1 |
| ***Bacillus* sp. ABP14** | **WP_070807265.1** | **NZ_CP017016.1** |
| ***Bacillus* sp. FDAARGOS_527** | **WP_063246657.1** | **NZ_CP033795.1** |
| ***Bacillus* sp. FJAT-42376** | **WP_123918462.1** | **NZ_CP033906.1** |
| ***Bacillus thuringiensis* serovar finitimus YBT-020** | **WP_000907289.1** | **NC_017200.1** |
| ***Bacillus weihenstephanensis* WSBC 10204** | **WP_033709612.1** | **NZ_CP009746.1** |
| *Bifidobacterium adolescentis* 22L | WP_038444142.1 | NZ_CP007443.1 |
| *Bifidobacterium adolescentis* 6 | WP_039774842.1 | NZ_CP023005.1 |
| ***Bifidobacterium adolescentis* ATCC 15703** | **WP_011742867.1** | **NC_008618.1** |
| *Bifidobacterium adolescentis* BBMN23 | WP_039774842.1 | NZ_CP010437.1 |
| *Bifidobacterium adolescentis* P2P3 | WP_039774842.1 | NZ_CP024959.1 |
| *Bifidobacterium breve* 017W439 | WP_052822180.1 | NZ_CP021554.1 |
| *Bifidobacterium breve* 082W48 | WP_106622134.1 | NZ_CP021555.1 |
| *Bifidobacterium breve* 12L | WP_019727643.1 | NZ_CP006711.1 |
| *Bifidobacterium breve* 139W423 | WP_021648898.1 | NZ_CP021556.1 |
| *Bifidobacterium breve* 180W83 | WP_106630814.1 | NZ_CP021557.1 |
| *Bifidobacterium breve* 215W447a | WP_106641225.1 | NZ_CP021558.1 |
| *Bifidobacterium breve* 689b | WP_025331928.1 | NZ_CP006715.1 |
| *Bifidobacterium breve* ACS-071-V-Sch8b | WP_014483227.1 | NC_017218.1 |
| *Bifidobacterium breve* BR3 | WP_052789711.1 | NZ_CP010413.1 |
| *Bifidobacterium breve* CNCM I-4321 | WP_106642177.1 | NZ_CP021559.1 |
| *Bifidobacterium breve* DRBB26 | WP_003833139.1 | NZ_CP021390.1 |
| *Bifidobacterium breve* DRBB27 | WP_106621239.1 | NZ_CP021552.1 |
| *Bifidobacterium breve* DRBB28 | WP_003833139.1 | NZ_CP021553.1 |
| *Bifidobacterium breve* DRBB29 | WP_106621239.1 | NZ_CP023198.1 |
| *Bifidobacterium breve* DRBB30 | WP_106642177.1 | NZ_CP023199.1 |
| ***Bifidobacterium breve* DSM 20213 = JCM 1192** | **WP_003827899.1** | **NZ_AP012324.1** |
| *Bifidobacterium breve* FDAARGOS_561 | WP_003827899.1 | NZ_CP033841.1 |
| *Bifidobacterium breve* lw01 | WP_016462023.1 | NZ_CP034192.1 |
| *Bifidobacterium breve* JCM 7017 | WP_025300916.1 | NZ_CP006712.1 |
| *Bifidobacterium breve* JCM 7019 | WP_025220954.1 | NZ_CP006713.1 |
| *Bifidobacterium breve* LMC520 | WP_077149301.1 | NZ_CP019596.1 |
| *Bifidobacterium breve* NCFB 2258 | WP_015438200.1 | NZ_CP006714.1 |
| *Bifidobacterium breve* NCTC 11815 | WP_003827899.1 | NZ_LR134348.1 |
| *Bifidobacterium breve* NRBB01 | WP_003827899.1 | NZ_CP021384.1 |
| *Bifidobacterium breve* NRBB02 | WP_106622507.1 | NZ_CP021385.1 |
| *Bifidobacterium breve* NRBB04 | WP_015438200.1 | NZ_CP021386.1 |
| *Bifidobacterium breve* NRBB08 | WP_106622507.1 | NZ_CP023192.1 |
| *Bifidobacterium breve* NRBB09 | WP_106629152.1 | NZ_CP021387.1 |
| *Bifidobacterium breve* NRBB11 | WP_106628600.1 | NZ_CP021388.1 |
| *Bifidobacterium breve* NRBB18 | WP_106622507.1 | NZ_CP023193.1 |
| *Bifidobacterium breve* NRBB19 | WP_106622507.1 | NZ_CP023194.1 |
| *Bifidobacterium breve* NRBB20 | WP_106622507.1 | NZ_CP023195.1 |
| *Bifidobacterium breve* NRBB27 | WP_106622507.1 | NZ_CP023196.1 |
| *Bifidobacterium breve* NRBB49 | WP_106622507.1 | NZ_CP023197.1 |
| *Bifidobacterium breve* NRBB50 | WP_106628600.1 | NZ_CP021391.1 |
| *Bifidobacterium breve* NRBB51 | WP_019727643.1 | NZ_CP021392.1 |
| *Bifidobacterium breve* NRBB52 | WP_080867748.1 | NZ_CP021393.1 |
| *Bifidobacterium breve* NRBB56 | WP_016462023.1 | NZ_CP021394.1 |
| *Bifidobacterium breve* NRBB57 | WP_052822180.1 | NZ_CP021389.1 |
| *Bifidobacterium breve* S27 | WP_025341481.1 | NZ_CP006716.1 |
| *Bifidobacterium breve* UCC2003 | WP_015438200.1 | NC_020517.1 |
| ***Bifidobacterium kashiwanohense* JCM 15439 = DSM 21854** | **WP_033500884.1** | **NZ_AP012327.1** |
| *Bifidobacterium kashiwanohense* PV20-2 | WP_039197257.1 | NZ_CP007456.1 |
| *Bifidobacterium longum* BXY01 | WP_013141166.1 | NZ_CP008885.1 |
| ***Bifidobacterium longum* subsp. infantis ATCC 15697 = JCM 1222 = DSM 20088** | **WP_014484651.1** | **NC_017219.1** |
| *Bifidobacterium longum* subsp. infantis BT1 | WP_060620711.1 | NZ_CP010411.1 |
| *Bifidobacterium longum* subsp. longum JDM301 | WP_013141166.1 | NC_014169.1 |
| ***Bifidobacterium pseudocatenulatum* DSM 20438 = JCM 1200 = LMG 10505** | **WP_004219882.1** | **NZ_AP012330.1** |
| *Brevibacillus brevis* DZQ7 | WP_048031022.1 | NZ_CP030117.1 |
| *Brevibacillus brevis* NBRC 100599 | WP_012684303.1 | NC_012491.1 |
| ***Brevibacillus brevis* NCTC 2611** | **WP_106652439.1** | **NZ_LR134338.1** |
| *Brevibacillus brevis* X23 | WP_017249674.1 | NZ_CP023474.1 |
| ***Brevibacillus formosus* NF2** | **WP_088910547.1** | **NZ_CP018145.1** |
| ***Butyrivibrio fibrisolvens* INBov1** | **WP_110073161.1** | **NZ_CM009896.1** |
| ***Butyrivibrio proteoclasticus* B316** | **WP_013281180.1** | **NC_014387.1** |
| *Clostridium botulinum* AM1195 | WP_061328351.1 | NZ_CP013701.1 |
| *Clostridium botulinum* CDC_67071 | WP_096043970.1 | NZ_CP013242.1 |
| *Clostridium botulinum* MAP 5 | WP_106899782.1 | NZ_CP027781.1 |
| *Clostridium botulinum* Mfbjulcb8 | WP_061328351.1 | NZ_CP027780.1 |
| ***Clostridium botulinum* Prevot_594** | **WP_040108159.1** | **NZ_CP006902.1** |
| *Clostridium perfringens* F262 | WP_003481250.1 | NZ_CM001477.1 |
| *Clostridium perfringens* FORC_003 | WP_003466749.1 | NZ_CP009557.1 |
| *Clostridium perfringens* FORC_025 | WP_070956832.1 | NZ_CP013101.1 |
| *Clostridium perfringens* JP838 | WP_061427915.1 | NZ_CP010994.1 |
| *Clostridium perfringens* JXJA17 | WP_003452556.1 | NZ_CP028149.1 |
| ***Clostridium perfringens* NCTC 2837** | **WP_003473144.1** | **NZ_LS483461.1** |
| ***Clostridium sporogenes* DSM 795** | **WP_003491425.1** | **NZ_CP011663.1** |
| *Clostridium sporogenes* NCIMB 10696 | WP_003491425.1 | NZ_CP009225.1 |
| ***Collinsella aerofaciens* C11** | **WP_117798150.1** | **NZ_CP024960.1** |
| *Collinsella aerofaciens* indica | WP_099431625.1 | NZ_CP024160.1 |
| ***Exiguobacterium oxidotolerans* N4-1P** | **WP_088837913.1** | **NZ_CP022236.1** |
| ***Exiguobacterium* sp. AT1b** | **WP_012727256.1** | **NC_012673.1** |
| ***Exiguobacterium* sp. MH3** | **WP_023469423.1** | **NC_022794.1** |
| ***Fictibacillus phosphorivorans* G25-29** | **WP_066398125.1** | **NZ_CP015378.1** |
| ***Halobacillus halophilus* DSM 2266** | **WP_014644340.1** | **NC_017668.1** |
| *Halobacillus halophilus* HL2HP6 | WP_014644340.1 | NZ_CP022106.1 |
| *Paenibacillus mucilaginosus* 3016 | WP_013920028.1 | NC_016935.1 |
| *Paenibacillus mucilaginosus* K02 | WP_013920028.1 | NC_017672.3 |
| ***Paenibacillus mucilaginosus* KNP414** | **WP_013920028.1** | **NC_015690.1** |
| ***Paenibacillu*s sp. BD3526** | **WP_060531382.1** | **NZ_CP013023.1** |
| ***Romboutsia* sp. Frifi** | **CEI72691.1** | **LN650648.1** |
| ***Roseburia hominis* A2-183** | **WP_014079902.1** | **NC_015977.1** |
| ***Roseburia intestinalis* M50/1** | **CBL09539.1** | **FP929049.1** |
| ***Rubrobacter xylanophilus* DSM 9941** | **WP_011565595.1** | **NC_008148.1** |
| ***Salimicrobium jeotgali* MJ3** | **WP_008592356.1** | **NZ_CP011361.2** |
| ***Selenomonas* sp. oral taxon 126 strain W7667** | **WP_066844117.1** | **NZ_CP016201.1** |
| ***Selenomona*s sp. oral taxon 478** | **WP_050342638.1** | **NZ_CP012071.1** |
| ***Selenomonas sputigena* ATCC 35185** | **WP_006193285.1** | **NC_015437.1** |
| *Streptococcus anginosus* C238 | WP_003035167.1 | NC_022239.1 |
| ***Streptococcus anginosus* NCTC 11169** | **WP_126407817.1** | **NZ_LR134288.1** |
| *Streptococcus anginosus* SA1 | WP_004224623.1 | NZ_CP007573.1 |
| ***Streptococcus equinus* FDAARGOS_251** | **WP_107373002.1** | **NZ_CP020438.1** |
| ***Streptococcus ferus* NCTC 12278** | **WP_018030127.1** | **NZ_LS483343.1** |
| *Streptococcus gallolyticus* NCTC 13773 | WP_077496569.1 | NZ_LS483409.1 |
| *Streptococcus gallolyticus* subsp. gallolyticus ATCC 43143 | WP_012961717.1 | NC_017576.1 |
| *Streptococcus gallolyticus* subsp. gallolyticus ATCC BAA-2069 | WP_013642905.1 | NC_015215.1 |
| ***Streptococcus gallolyticus* subsp. gallolyticus DSM 16831** | **WP_077496569.1** | **NZ_CP018822.1** |
| *Streptococcus gallolyticus* UCN34 | WP_012961717.1 | NC_013798.1 |
| *Streptococcus gordonii* FDAARGOS_371 | WP_061595675.1 | NZ_CP023511.1 |
| *Streptococcus gordonii* IE35 | WP_046165000.1 | NZ_CP017295.1 |
| *Streptococcus gordonii* KCOM 1506 (= ChDC B679) | WP_053794170.1 | NZ_CP012648.1 |
| *Streptococcus gordonii* NCTC 3165 | WP_111723252.1 | NZ_LS483375.1 |
| ***Streptococcus gordonii* NCTC 7865** | **WP_060553094.1** | **NZ_LS483341.1** |
| *Streptococcus gordonii* NCTC 7868 | WP_011999856.1 | NZ_LR134291.1 |
| *Streptococcus gordonii* str. Challis substr. CH1 | WP_011999856.1 | NC_009785.1 |
| *Streptococcus intermedius* B196 | WP_021003212.1 | NC_022246.1 |
| *Streptococcus intermedius* C270 | WP_003076142.1 | NC_022237.1 |
| *Streptococcus intermedius* FDAARGOS_233 | WP_082312092.1 | NZ_CP020433.2 |
| ***Streptococcus intermedius* JTH08** | **WP_003076142.1** | **NC_018073.1** |
| *Streptococcus intermedius* NCTC 11324 | WP_003076142.1 | NZ_LS483436.1 |
| *Streptococcus intermedius* TYG1620 | WP_003076142.1 | NZ_AP014880.1 |
| ***Streptococcus pantholopis* TA 26** | **WP_067060972.1** | **NZ_CP014699.1** |
| ***Streptococcus pasteurianus* ATCC 43144** | **WP_003063947.1** | **NC_015600.1** |
| *Streptococcus pasteurianus* NCTC 13784 | WP_111718197.1 | NZ_LS483462.1 |
| *Streptococcus salivarius* ATCC 27945 | WP_084914613.1 | NZ_CP015282.1 |
| *Streptococcus salivarius* HSISS4 | WP_021144275.1 | NZ_CP013216.1 |
| *Streptococcus salivarius* JF | WP_022496455.1 | NZ_CP014144.1 |
| ***Streptococcus salivarius* JIM8777** | **WP_014634402.1** | **NC_017595.1** |
| *Streptococcus salivarius* NCTC 8618 | WP_022496455.1 | NZ_CP009913.1 |
| *Streptococcus* sp. FDAARGOS_192 | WP_080610467.1 | NZ_CP020431.2 |
| ***Streptococcus* sp. HSISS1** | **EQC64892.1** | **CM002132.1** |
| ***Streptococcus* sp. Z15** | **WP_116877779.1** | **NZ_CP031733.1** |
| *Streptococcus thermophilus* APC151 | WP_014608381.1 | NZ_CP019935.1 |
| ***Streptococcus thermophilus* JIM 8232** | **WP_014621674.1** | **NC_017581.1** |
| *Streptococcus thermophilus* LMD-9 | ABJ66323.1 | CP000419.1 |
| *Streptococcus thermophilus* MN-BM-A01 | WP_014727436.1 | NZ_CP012588.1 |
| *Streptococcus thermophilus* MN-ZLW-002 | WP_014727436.1 | NC_017927.1 |
| *Streptococcus thermophilus* MTH17CL396 | ETW89989.1 | CM002371.1 |
| *Streptococcus thermophilus* ND03 | WP_014608381.1 | NC_017563.1 |
| *Streptococcus thermophilus* SMQ-301 | WP_046206469.1 | NZ_CP011217.1 |
| *Streptococcus thermophilus* TH1477 | WP_071417354.1 | NZ_CM003135.1 |
| ***Tumebacillus algifaecis* THMBR28** | **WP_094236481.1** | **NZ_CP022657.1** |
| ***Tumebacillus avium* AR23208** | **WP_087458314.1** | **NZ_CP021434.1** |

* Species listed in bold are represented in Fig. 1.

**Table S2: Bacterial strains and plasmids used in this study**

| **Strain** | **Description*** | **Source** | |
| --- | --- | --- | --- |
| ***E. coli*** | | | |
| DH5α | Cloning strain; F^–^ Φ80*lacZ*ΔM15 Δ(*lacZYA*-*argF*) U169 *recA1* *endA1 hsdR17* (r_K_^–^, m_K_^+^) *phoA supE44* λ^–^ *thi*-1 *gyrA96* *relA1* | Invitrogen | |
| EC135 | Strain lacking endogenous restriction modification systems and DNA methyltransferases; TOP10 Δ*dam* Δ*dcm* Δ*hsd* Δ*mcrBC* Δ*mcrA* Δ*mrr* | [1] | |
| BL21CodonPlus^TM^  (DE3)-RP | Protein expression strain; F^-^ *ompT* *hsdS*(rB^−^ mB^−^) *dcm*^+^ Tet^r^ *gal*λ (DE3) *end*AHte *met*A∷Tn5(Kan^r^) [*arg*U *ile*Y *leu*W Cam^r^] | Stratagene | |
| ***B. cereus*** | | | |
| ATCC 10987 | Wild-type strain | A.J. Clarke | |
| ATCC 10987 Δ*pelA_H_* | ATCC 10987 with an unmarked, non-polar deletion of *pelA_H_* (*BCE_5582*) | This study | |
| ATCC 10987 Δ*pelD* | ATCC 10987 with an unmarked, non-polar deletion of *pelD* (*BCE_5583*) | This study | |
| ATCC 10987 Δ*pelE* | ATCC 10987 with an unmarked, non-polar deletion of *pelE* (*BCE_5584*) | This study | |
| ATCC 10987 Δ*pelA_DA_* | ATCC 10987 with an unmarked, non-polar deletion of *pelA_DA_* (*BCE_5585*) | This study | |
| ATCC 10987 Δ*pelF* | ATCC 10987 with an unmarked, non-polar deletion of *pelF* (*BCE_5586*) | This study | |
| ATCC 10987 Δ*pelG* | ATCC 10987 with an unmarked, non-polar deletion of *pelG* (*BCE_5587*) | This study | |
| ATCC 10987 Δ*cdgF* | ATCC 10987 with an unmarked, non-polar deletion of *cdgF* (*BCE_0696*) | This study | |
| ATCC 10987 Δ*cdgE* | ATCC 10987 with an unmarked, non-polar deletion of *cdgE* (*BCE_3781*) | This study | |
| ATCC 10987 *pelD*^R363A^ | ATCC 10987 with a mutation of arginine 363 encoded by *pelD* (*BCE_5583*) to alanine | This study | |
| ATCC 10987 *pelD*^D366A^ | ATCC 10987 with a mutation of aspartic acid 366 encoded by *pelD* (*BCE_5583*) to alanine | This study | |
| ATCC 10987 *pelD*^R395A^ | ATCC 10987 with a mutation of arginine 395 encoded by *pelD* (*BCE_5583*) to alanine | This study | |
| **Plasmid** | **Description** | **Source** | |
| **Recombinant protein expression** | | | |
| pM.Bce | Arabinose inducible vector to express *B. cereus* ATCC 10987 DNA methyltransferases; methylates co-replicating plasmids to overcome the restriction barrier of ATCC 10987; Spc^R^ | | [1] |
| pET-24a(+) | IPTG inducible protein expression vector encoding a C-terminal hexahistidine tag, Kan^R^ | | Novagen |
| pET-28a(+) | IPTG inducible protein expression vector encoding an N-terminal hexahistidine tag and thrombin cleavage site, Kan^R^ | | Novagen |
| pET24a::PelA_H_*^Bc^* | *B. cereus* ATCC 10987 PelA_H_ (BCE_5582), excluding the predicted signal sequence (residues 1-21), inserted between the BamHI and XhoI sites of pET-24a(+) | | This study |
| pET24a::PelA_H_*^Bc-^*^E213A^ | pET24a::PelA_H_*^Bc^* with a mutation of glutamate 213 to alanine | | This study |
| pET28a::PelA_47-303_ | *P. aeruginosa* PAO1 *pelA*, encoding the glycoside hydrolase domain (residues 47-303), inserted between the NdeI and XhoI sites of pET-28a(+) | | [2] |
| pET28a::PelA_47-303_^E218A^ | pET28a::PelA_47-303_ with a mutation of glutamate 218 to alanine | | [2] |
| pET28a::PelD_150-407_ | *B. cereus* ATCC 10987 *pelD* (*BCE_5583*), encoding the predicted cytoplasmic domain of the protein (residues 150-407), inserted between the NheI and XhoI sites of pET-28a(+) | | This study |
| pET28a::PelD_150-407_^R363A^ | pET28a::PelD_150-407_ with a mutation of arginine 363 to alanine | | This study |
| pET28a::PelD_150-407_^D366A^ | pET28a::PelD_150-407_ with a mutation of aspartic acid 366 to alanine | | This study |
| pET28a::PelD_150-407_^R395A^ | pET28a::PelD_150-407_ with a mutation of arginine 395 to alanine | | This study |
| **Allelic exchange** | | | |
| pMAD | *E. coli* – *Bacillus* shuttle vector, encoding the thermostable β-galactosidase *bgaB* from *Bacillus stearothermophilus* driven by the constitutive P*clpB* promoter; Erm^R^, Amp^R^ | | [3] |
| pMAD::Δ*pelA_H_* | *B. cereus* ATCC 10987 Δ*pelA_H_* (*BCE_5582*) cloned between the BamHI and NcoI sites of pMAD | | This study |
| pMAD::Δ*pelD* | *B. cereus* ATCC 10987 Δ*pelD* (Δ*BCE_5583*) cloned between the BamHI and SmaI sites of pMAD | | This study |
| pMAD::Δ*pelE* | *B. cereus* ATCC 10987 Δ*pelE* (Δ*BCE_5584*) cloned between the BamHI and SmaI sites of pMAD | | This study |
| pMAD::Δ*pelA_DA_* | *B. cereus* ATCC 10987 Δ*pelA_DA_* (Δ*BCE_5585*) cloned between the BamHI and SmaI sites of pMAD | | This study |
| pMAD::Δ*pelF* | *B. cereus* ATCC 10987 Δ*pelF* (Δ*BCE_5586*) cloned between the KpnI and BamHI sites of pMAD | | This study |
| pMAD::Δ*pelG* | *B. cereus* ATCC 10987 Δ*pelG* (Δ*BCE_5587*) cloned between the SalI and SmaI sites of pMAD | | This study |
| pMAD::Δ*cdgF* | *B. cereus* ATCC 10987 Δ*cdgF* (Δ*BCE_0696*) cloned between the BamHI and SmaI sites of pMAD | | This study |
| pMAD::Δ*cdgE* | *B. cereus* ATCC 10987 Δ*cdgE* (Δ*BCE_3781*) cloned between the SalI and SmaI sites of pMAD | | This study |
| pMAD::*pelD*^R363A^ | *B. cereus* ATCC 10987 *pelD* (*BCE_5583*), with a mutation of arginine 363 to alanine, cloned between the BamHI and EcoRI sites of pMAD | | This study |
| pMAD::*pelD*^D366A^ | *B. cereus* ATCC 10987 *pelD* (*BCE_5583*), with a mutation of aspartic acid 366 to alanine, cloned between the BamHI and EcoRI sites of pMAD | | This study |
| pMAD::*pelD*^R395A^ | *B. cereus* ATCC 10987 *pelD* (*BCE_5583*), with a mutation of arginine 395 to alanine, cloned between the BamHI and EcoRI sites of pMAD | | This study |
| **Complementation in *B. cereus*** | | | |
| pAD123 | *E. coli* – *Bacillus cereus* shuttle vector, encodes a promoterless copy of GFPmut3a for promoter screening; Cam^R^, Amp^R^ | | [4] |
| pHCMC04 | *E. coli* – *Bacillus subtilis* shuttle vector, contains the xylose-inducible *xylR*-P*_xylA_* promoter cassette; Cam^R^, Amp^R^ | | [5] |
| pAD123-P_xyl_ | pAD123 with the *xylR*-P*_xylA_* cassette from pHCMC04 cloned between the SacI and BamHI sites of pAD123; contains a multiple cloning site (EcoRV-KpnI-NheI-NotI-SmaI-BamHI) immediately downstream of P*_xylA_* | | This study |
| pAD123-P_xyl_::*pelA_H_* | *B. cereus* ATCC 10987 *pelA_H_* (*BCE_5582*) fused to a synthetic RBS (5’-TAAGGAGGAAGCAGGT-3’) cloned between the EcoRV and BamHI sites of pAD123-P_xyl_ | | This study |
| pAD123-P_xyl_::*pelD* | *B. cereus* ATCC 10987 *pelD* (*BCE_5583*) fused to a synthetic RBS (5’-TAAGGAGGAAGCAGGT-3’) cloned between the KpnI and BamHI sites of pAD123-P_xyl_ | | This study |
| pAD123-P_xyl_::*pelE* | *B. cereus* ATCC 10987 *pelE* (*BCE_5584*) fused to a synthetic RBS (5’-TAAGGAGGAAGCAGGT-3’) cloned between the EcoRV and BamHI sites of pAD123-P_xyl_ | | This study |
| pAD123-P_xyl_::*pelA_DA_* | *B. cereus* ATCC 10987 *pelA_DA_* (*BCE_5585*) fused to a synthetic RBS (5’-TAAGGAGGAAGCAGGT-3’) cloned between the KpnI and BamHI sites of pAD123-P_xyl_ | | This study |
| pAD123-P_xyl_::*pelF* | *B. cereus* ATCC 10987 *pelF* (*BCE_5586*) fused to a synthetic RBS (5’-TAAGGAGGAAGCAGGT-3’) cloned between the KpnI and BamHI sites of pAD123-P_xyl_ | | This study |
| pAD123-P_xyl_::*pelG* | *B. cereus* ATCC 10987 *pelG* (*BCE_5587*) fused to a synthetic RBS (5’-TAAGGAGGAAGCAGGT-3’) cloned between the EcoRV and SmaI sites of pAD123-P_xyl_ | | This study |
| pAD123-P_xyl_::*cdgF* | *B. cereus* ATCC 10987 *cdgF* (*BCE_0696*) fused to a synthetic RBS (5’-TAAGGAGGAAGCAGGT-3’) cloned between the KpnI and BamHI sites of pAD123-P_xyl_ | | This study |
| pAD123-P_xyl_::*cdgE* | *B. cereus* ATCC 10987 *cdgE* (*BCE_3781*) fused to a synthetic RBS (5’-TAAGGAGGAAGCAGGT-3’) cloned between the EcoRV and BamHI sites of pAD123-P_xyl_ | | This study |

*Amp, ampicillin; Cam, chloramphenicol; Kan, kanamycin; Erm, erythromycin; Spc, spectinomycin

**Table S3: Primers used in this study**

| **Primers** | **Sequence*** |
| --- | --- |
| **Vectors for recombinant protein production** | |
| PelA_H_*^Bc^* pET-F | GGG **GGA TCC** AAT GTC GTG GAA CCA GTT TTA AAA |
| PelA_H_*^Bc^* pET-R | GGG **CTC GAG** TTT ATC ATA AAT ATC ATT CGG ATT AAT TG |
| PelA_H_*^Bc^* E213A upR | T ATT GAA GTC CgC CCA CAA TAC A |
| PelA_H_*^Bc^* E213A downF | T GTA TTG TGG GcG GAC TTC AAT A |
| PelD_Bc_ 150F | CG **GCT AGC** GAA TCA AAT AAT CAC TTA TCT AAA ATG TAT C |
| PelD_Bc_ 407R | GC **CTC GAG** TTA GCA TAC CAC TCC TTT AGA AGA AA |
| PelD_Bc_ R363A F | AAA GAT CAT TTG gcT GAT ATC GAT ATA |
| PelD_Bc_ R363A R | TAT ATC GAT ATC Agc CAA ATG ATC TTT |
| PelD_Bc_ D366A F | TTG CGT GAT ATC GcT ATA TTC GGT TA |
| PelD_Bc_ D366A R | TA ACC GAA TAT AgC GAT ATC ACG CAA |
| PelD_Bc_ R395A F | A GTC CAA ACT gcT ATA CAA AAC GC |
| PelD_Bc_ R395A R | GC GTT TTG TAT Agc AGT TTG GAC T |
| **Allelic exchange vectors** |  |
| ΔpelA_H_ upF | GTG **GGA TCC** GTA TTC GAT CTG TTT TTC GTA CCG |
| ΔpelA_H_ upR | *ATA AAT ATC ATT CGG ATT AAT TGT AGG* AAA GTA CAT GGA ACG TTT CCA CTC |
| ΔpelA_H_ downF | CCT ACA ATT AAT CCG AAT GAT ATT TAT |
| ΔpelA_H_ downR | GGA **CCA TGG** CAA CCC TCG GAA CGG GAA TC |
| ΔpelD upF | GGT **GGA TCC** CTA AAT AGG TCA GAC TTC CTA CC |
| ΔpelD upR | *TTA GCA TAC CAC TCC TTT AGA* TAA ATT AGA TTG TCT ATA GTG CAT |
| ΔpelD downF | TCT AAA GGA GTG GTA TGC TAA |
| ΔpelD downR | GAG **CCC GGG** AGT TGT ATT CTC ATC TAG CGT AAT AT |
| ΔpelE upF | GGG **GGA TCC** GAT GAT ATAT TAC TGT ACT TTT ATT CAT TAG |
| ΔpelE upR | *AAT GTG TGC GCC TCC AAA CTG* AAA GTA AAG AAG GAG TAA AAT CAT |
| ΔpelE downF | CAG TTT GGA GGC GCA CAC ATT |
| ΔpelE downR | GTG **CCC GGG** TTC TCT CCC AAA TAA TAA ATA GGT ATT |
| ΔpelA_DA_ upF | GAG **GGA TCC** ACT GGT AAA AAT ACC TTA CCT CAA TC |
| ΔpelA_DA_ upR | *TAC CCC TTT AAT TGG AAT TGT AGC* GAT ATA TAT ATT AAT CCC CTT TTT CAA |
| ΔpelA_DA_ downF | GCT ACA ATT CCA ATT AAA GGG GTA |
| ΔpelA_DA_ downR | GTG **CCC GGG** CTT TTA CCC TTG AAG AAT AGT AGT CT |
| ΔpelF upF | TAG TAC AGA **GTC GAC** GGA AGT AAA AGT AAC GGC AGT TTA AT |
| ΔpelF upR | *ATA CCT GCC ATG TTT TCG CAC C*GC CAC TGA CAA AAG GAT AAC TTC C |
| ΔpelF downF | GGT GCG AAA ACA TGG CAG GTA T |
| ΔpelF downR | TAG TAC AGA **CCC GGG** GAT CCG ATT AAA AAT GCC AGT GCA |
| ΔpelG upF | ACG **GTC GAC** GCC AAA ACA TAA TTC AGA TAA AC |
| ΔpelG upR | *TTA TTT CCC GTT AAG TTT ATT AGC* TCG AAA TCC TAT ACC TGC CAT |
| ΔpelG downF | GCT AAT AAA CTT AAC GGG AAA TAA |
| ΔpelG downR | TAT **CCC GGG** CAG ATA GAG AAT GGA GTA TCG |
| ΔcdgF upF | AGG **GGA TCC** TTC TTC TGA ACG ATG GAG ATGC |
| ΔcdgF upR | *TAA CTT TTC TTT TAA CTC TCC TAC AGA* TGC ATG ACC TCT TTG TTC TAG CAT |
| ΔcdgF downF | TCT GTA GGA GAG TTA AAA GAA AAG TTA |
| ΔcdgF downR | GGT **CCC GGG** GTT TTA ACA GCT TTT GTT ATC GGG |
| ΔcdgE upF | GGG **GTC GAC** TAT TTC TGT TAC TCT ATG AGC GTC T |
| ΔcdgE upR | *GAG AAG AGT AGA AAA TGA ATC AGC* TGT CGC AAT TAA TAT ACA TAG TTG TAA |
| ΔcdgE downF | GCT GAT TCA TTT TCT ACT CTT CTC |
| ΔcdgE downR | GAG **CCC GGG** TGA AGG GAT CGT GGA TAT AAG C |
| pelD I-site F | GGC **GGA TCC** CGA AAG AAG GGA TTT TTT ATG CAC TA |
| pelD I-site R | GCC **GAA TTC** CTT GTT GCC CTA ATG TAT TAT CCG |
| pelD R363A upR | A TAT ATC GAT ATC Agc CAA ATG ATC TTT TAA AAT AGA ATT TAT CAT TTC TAA AGA A |
| pelD R363A downF | A GAT CAT TTG gcT GAT ATC GAT ATA TTC GGT TAT AGC ACG ACT AAA CAA |
| pelD D366A upR | ATA TAG CgA TAT CAC GCA AAT GAT CTT TTA AAA TAG AAT TTA TCA TTT CTA AAG AA |
| pelD D366A downF | A GAT CAT TTG CGT GAT ATC GcT ATA TTC GGT TAT AGC ACG ACT AAA CAA |
| pelD R395A upR | C GTT TTG TAT Agc AGT TTG GAC TGG TAA TAA AAA CTT TTC TTC AGT GCC A |
| pelD R395A downF | CA GTC CAA ACT gcT ATA CAA AAC GCT CTT TCT TCT AAA GGA GTG GTA |
| **Complementation vectors** | |
| xylR-PxylA F | GGG **GAG CTC** CTA ACT TAT AGG GGT AAC ACT TA |
| xylR-PxylA R | GAA **GGA TCC** CAT TTC CCC CTT TGA TTT TTA GA |
| Pxyl MCS F | AG CGC GGC CGC CCC GGG GGA TCC TCT AGA TTT AAG AAG GAG |
| Pxyl MCS R | AGC GGT ACC GAT ATC CAT TTC CCC CTT TGA TTT AAG TGA A |
| pelA_H_ F | GGT **GAT ATC *TAA GGA GGA AGC AGG T***AT GGA GTG GAA ACG TTC CAT GTA |
| pelA_H_ R | GGT **GGA TCC** CTA TTT ATC ATA AAT ATC ATT CGG ATT AA |
| pelD F | GAG **GGT ACC *TAA GGA GGA AGC AGG T***AT GCA CTA TAG ACA ATC TAA TTT ATT AC |
| pelD R | GAG **GGA TCC** TTA GCA TAC CAC TCC TTT AGA AGA |
| pelE F | GGG **GAT ATC *TAA GGA GGA AGC AGG T***AT GAT TTT ACT CCT TCT TTA CTT TAT T |
| pelE R | GTG **GGA TCC** TTA ATC CCC TTT TTC AAT GTG TGC |
| pelA_DA_ F | GAG **GGT ACC *TAA GGA GGA AGC AGG T***TT GAA AAA GGG GAT TAA TAT ATA TAT CG |
| pelA_DA_ R | GTG **GGA TCC** CTA TAC CCC TTT AAT TGG AAT TGT AG |
| pelF F | GAG **GGT ACC *TAA GGA GGA AGC AGG T***AT GAG AAT AGG TTT AGT CGT TGA AG |
| pelF R | GGT **GGA TCC** CTA TAC CTG CCA TGT TTT CGC A |
| pelG F | GGG **GAT ATC *TAA GGA GGA AGC AGG T***AT GGC AGG TAT AGG ATT TCG ATT A |
| pelG R | GAG **CCC GGG** TTA TTT CCC GTT AAG TTT ATT AGC TAA |
| cdgF F | GAG **GGT ACC *TAA GGA GGA AGC AGG T***AT GCT AGA ACA AAG AGG TCA TGC A |
| cdgF R | GTG **GGA TCC** TTA TAT GAA ATC TGT AGT TAA CTT TTC |
| cdgE F | GGT **GAT ATC *TAA GGA GGA AGC AGG T***GT GAT TTT ATT GAA ATT AAA TAA AAA T |
| cdgE R | GGT **GGA TCC** CTA CAG AGA ATG AAC ACC TTT T |
| **Sequencing primers** | |
| T7 | TAA TAC GAC TCA CTA TAG GG |
| T7ter | GCT AGT TAT TGC TCA GCG G |
| pMAD SEQ-F | GCA ACG CGG GCA TCC CGA TG |
| pMAD SEQ-R | CCC AAT ATA ATC ATT TAT CAA CTC TTT TAC ACT TAA ATT TCC |
| pAD123-P_xyl_ SEQ-F | GAT AGT TGA TGG ATA AAC TTG TTC |
| pAD123-P_xyl_ SEQ-R | CAA GAA TTG GGA CAA CTC CAG |
| pelA_DA_ SEQ-int | GTT CAA TCA CTG GGT ACT ATT TTC |
| cdgF SEQ-int | ATG CTT TCA GGC ACC ATT TAT GC |
| cdgE SEQ-int-1 | CTG AAA CGA TAA CAT TAT TAC GTC |
| cdgE SEQ-int-2 | GAA AAT TGG GAA CCG TGC CTT T |
| cdgE SEQ-int-3 | CAA GTT GAT AGC GAA TCT AAC GAC |

*Restriction sites are bolded; regions of complementary to the target amplicon are underlined; regions of reverse complementarity (to facilitate splicing) are italicized; lowercase letters denote a nucleotide substitution; synthetic ribosomal binding sites are in bold italics.
